# IL-15 superagonist induces robust STAT5 phosphorylation to support the differentiation and survival of tumor-relevant memory precursor effector-like CD8 T cells, enhancing long-term antitumor protection

**DOI:** 10.1101/2025.08.12.669959

**Authors:** Tatiana Delgado Cruz, Rachel Honigsberg, Geoffrey J. Markowitz, Bharati Swami, Arshdeep Singh, Meixian Stephanie Tang, Liron Yoffe, Sally E. Claridge, Stella A. Martomo, Grayson E. Rodriguez, K. Christopher Garcia, Yongfeng He, Yang Bai, Jeegar Patel, Matthew Pang, Dicle Özge Özgenel, Marissa M. Michael, Juan Cubillos-Ruiz, Timothy E. McGraw, Olivier Elemento, Nasser K. Altorki, Vivek Mittal, Jonathan Villena-Vargas

## Abstract

**Background:** Common gamma chain cytokines drive CD8 T cells to proliferate and acquire effector function, but strong interleukin-2 (IL-2) signaling also drives terminal differentiation at the expense of memory. Engineered cytokine agonists are now entering the clinic, yet which property of cytokine signaling determines whether an expanding effector retains memory potential and the role of memory in long term responses remains unresolved.

**Methods:** We use a resectable, spontaneously metastatic mouse model of lung adenocarcinoma to compare an IL-15 superagonist with PD-1 blockade, alone and in combination, and to follow tumor-relevant CD8 T-cell clones from treatment through surgery to metastatic relapse. We combine flow cytometry with paired single-cell RNA and T-cell receptor sequencing of tumor, tumor-draining and non-draining lymph nodes. We use receptor blockade and phosphorylation flow cytometry in OT-1 cultures to define the signaling requirement, and subset depletion with tumor rechallenge to test the resulting memory.

**Results:** The IL-15 superagonist expands a cytotoxic CD8 T-cell state that retains IL-7Rα and lacks KLRG1, which we term memory-precursor effector-like (Tmpe). This state requires CD122 and accompanies STAT5 phosphorylation exceeding that induced by conventional IL-15 or IL-2, with minimal STAT3 engagement. Tumor-matched clonal diversity increases within a progenitor-like compartment of the tumor-draining lymph node, and the clones carrying this program are those recovered in non-draining lymph nodes long after tumor resection. Combination with PD-1 blockade reduces metastatic recurrence and extends survival. Unexpectedly, depletion of either CD4 or CD8 T cells alone fails to abrogate protection on rechallenge and only combined depletion does, unlike the memory established by PD-1 blockade, where loss of either subset is sufficient.

**Conclusions:** Our data link the Tmpe state to CD122-dependent signaling accompanied by strong STAT5 and minimal STAT3 phosphorylation, and show that the clones carrying this state are those that persist in lymphoid tissue and protect against recurrence. This suggests that how an engineered cytokine engages STAT proteins, and not only how strongly, shapes whether an expanding effector retains memory potential, which is a testable design parameter for the next generation of cytokine agonists.

**Key Message:** *What is already known on this topic:* Strong γc cytokine signaling, exemplified by IL-2, drives CD8 T cells toward terminal effector differentiation at the expense of memory, and engineered cytokine agonists have entered the clinic without a defined principle for which signaling property should be optimized.

*What this study adds:* An IL-15 superagonist that drives strong STAT5 phosphorylation with minimal STAT3 engagement generates a CD122-dependent cytotoxic CD8 T cell state that retains IL-7Rα, with associated persistence in lymphoid tissue after tumor resection and protect against metastatic recurrence. This memory is functionally distinct, since either CD4 or CD8 T cells alone are sufficient for protection on rechallenge, unlike the memory established by PD-1 blockade.

*How this study might affect research, practice or policy:* These findings indicate that how an engineered cytokine engages STAT proteins as a design parameter for durable antitumor immunity, and that the durability of a cytokine agonist is best assessed in models permitting clonal tracking through resection into relapse.

## Background

Harnessing the capacity of T cells to specifically identify and destroy tumor cells through immune checkpoint blockade (ICB) therapy has become a major therapeutic breakthrough in many types of cancer, including lung cancer, moving from later-line treatment options to first-line therapies [1]. Increasing clinical and preclinical evidence has shown that PD-1/PD-L1 immune checkpoint blockade not only promotes antitumor T cell responses by reinvigorating highly differentiated effector CD8^+^ T cells in the primary tumor, but also relies on other less differentiated subsets, such as progenitor exhausted T cells (Tpex), as exhausted T cells in the tumor possess limited proliferative and functional capacity [2, 3].

Recent evidence indicates that PD-1 ICB relies on Tpex that retain polyfunctionality, can persist long-term, as well as differentiate into terminally exhausted T cells [4-6]. Tumor-draining lymph nodes (tdLN) are enriched in Tpex, which proliferate at a high rate after ICB treatment [7, 8], with data suggesting that these progenitors migrate from the tdLN to the tumor after therapy to enhance antitumor immunity [9-11]. The abundance of this population has been correlated with response to ICB. Consequently, therapeutic strategies that enhance this specific T cell population may yield superior tumor control and improve clinical outcomes [12, 13].

Most patients with non-small cell lung cancer (NSCLC) do not respond to anti-PD-1 (aPD-1) therapy, either alone or in combination with chemotherapy [14], prompting strong interest in novel combination strategies to enhance ICB efficacy.

Cytokine-based therapies have been explored in cancer treatment for decades. Among these, IL-2 has been extensively studied as a T cell growth factor that provides strong stimulatory signals for anti-tumor activity by supporting proliferation, activation, and differentiation into cytolytic effector T cells [15] [16].

High-dose recombinant IL-2 therapy has previously shown remarkable responses in patients with metastatic melanoma, even achieving complete responses in a subset of patients [17]. Unfortunately the implementation of cytokine based therapies at the clinical level has been historically limited by its narrow therapeutic index[18]. The development of novel molecules with higher potency and improved therapeutic prolifes has made it feasible to reconsider their clinical implementation.

More recently, Hashimoto et al. showed that IL-2 alone or in combination with aPD-1 results in downregulation of several inhibitory receptors, as well as upregulation of effector molecules and inflammatory cytokines, suggesting that this cytokine can modulate exhaustion programs to generate highly functional CD8^+^ T cells [19, 20]. IL-15 and IL-2 belong to the same family of cytokines and share two of their three receptor subunits (CD122 and CD132), but differ in their respective α chains (CD215 and CD25, respectively), resulting in overlapping biological functions [21]. Despite these similarities, IL-15 uniquely supports memory formation and persistence of CD8^+^ T cells [16, 22] and promotes self-renewal of precursor exhausted CD8^+^ T cell subsets rather than rescuing terminally differentiated effectors [23-25], suggesting that IL-15 may provide greater antitumor control and promote lasting T cell memory.

In this work, we used an immunocompetent mouse model of lung adenocarcinoma with a highly metastatic lung cancer cell line to understand the effect of an IL-15 superagonist on T cell responses in the acute setting, as well as its impact on tumor progression, metastasis, and long-term memory. Here, we show that IL-15 superagonist resulted in robust tumor control and modulation of the exhaustion program of tumor-relevant CD8^+^ T cells at the tumor and tdLN level, resulting in the generation of a highly functional memory-precursor effector-like T cell (Tmpe), an effector subset characterized by IL-7Rα^+^, KLRG1^-^ expression and cytotoxic features that retains plasticity to differentiate into other memory subsets [26]. IL-15 superagonism also increased the proportion of tumor-relevant clones observed at the memory stage, resulting in overall superior primary antitumor control, higher persistence of tumor memory-like T cell clones, and decreased metastatic recurrence when compared with aPD-1 alone. Mechanistically, this phenotype was associated with robust STAT5 phosphorylation compared with conventional IL-15 or IL-2. These findings provide a strong rationale for the use of an IL-15 superagonist and other novel non-natural cytokines that rewire STAT signaling to modulate T cell responses to enhance antitumor immunity beyond that achieved with standard ICB.

## RESULTS

### IL-15 superagonist results in robust tumor control and induces a memory-precursor effector-like T cell (Tmpe) phenotype *in vivo*

Proliferation of progenitor-like TCF1^+^PD-1^+^CD8^+^ T cells or Tpex cells is critical for the efficacy of PD-1 blockade [5, 12, 27]. Combination therapies to achieve durable responses have shown promising results, particularly the use of cytokines that provide growth signals for T cells [28]. IL-15 is integral to T cell survival [29], and has also been shown to promote self-renewal of Tpex cells [23].

We hypothesized that combining neoadjuvant PD-1 blockade with an IL-15 agonist would potentiate antitumor immunity by promoting CD8^+^ Tpex T cells in the tumor and tdLN [30-33]. To test this hypothesis, we utilized the IL-15 superagonist KD033, a fusion immunocytokine that comprises a high-affinity anti-PD-L1 antibody linked to the IL-15/IL-15Rα sushi domain complex. This design facilitates IL-15 trans-presentation from PD-L1-expressing immune cells and tumor cells in the tumor microenvironment (TME), increasing cytokine half-life and bioavailability while minimizing antibody-dependent cellular cytotoxicity (ADCC) [34].

Characterization of KD033 showed that its antitumor efficacy in a colon carcinoma mouse model relied on CD8^+^ T cells with a unique phenotype characterized by upregulation of genes associated with cytotoxicity, in contrast with untreated controls [34]. It was also shown that the IL-15 component of KD033 was responsible for most of this phenotype, as changes in gene signatures induced by this superagonist were recapitulated in a non-targeting variant (nt-KD033) comprising only the IL-15/15Rα fraction [34].

To assess the antitumor efficacy of IL-15 superagonism and its effect on metastatic recurrence, we employed a murine model of metastatic NSCLC using the highly metastatic 344SQ (*Kras^G12D^ p53^R172H^*^Δ*G*^) cell line (**Supplementary Fig. 1a-f**) [35]. Primary tumors were established in the right flank of 129Sv mice, enabling subsequent resection by survival surgery for evaluation of distant metastatic disease, memory formation, and survival. The tdLN (right inguinal lymph node) and non-tumor draining lymph node (ntdLN), or contralateral inguinal node, in this model were defined by their anatomical proximity to the tumor (right flank) [36]. Additionally, flow cytometric analyses revealed an increased percentage of PD-1^+^, interferon-γ, and Ki67-expressing CD8^+^ T cells in the tdLN relative to ntdLN (**Supplementary Fig. 1g, h)**, supporting the notion of a localized, nodal immune response.

Administration of a single dose of IL-15 superagonist, either alone or in combination with three doses of aPD-1, significantly reduced primary tumor burden compared with aPD-1 monotherapy or isotype control. By days 7-10 post-treatment, >90% of mice receiving IL-15 superagonist exhibited reduced BLI signal relative to control mice, compared with 30-50% under PD-1 blockade alone (**Fig. 1a, b**). Initial evaluation of immune populations at the tdLN by flow cytometry showed that IL-15 superagonist either alone or in combination with aPD-1 resulted in a significant proportional expansion of CD8^+^ T cells compared with aPD-1 alone and control groups from approximately 20% to 50% with a similar trend at the tumor level **(Supplementary Fig. 1i, j)**, suggesting that IL-15 superagonist was inducing phenotypic changes in the CD8^+^ T cell compartment that may contribute to the robust antitumor control observed.

**Figure 1.**
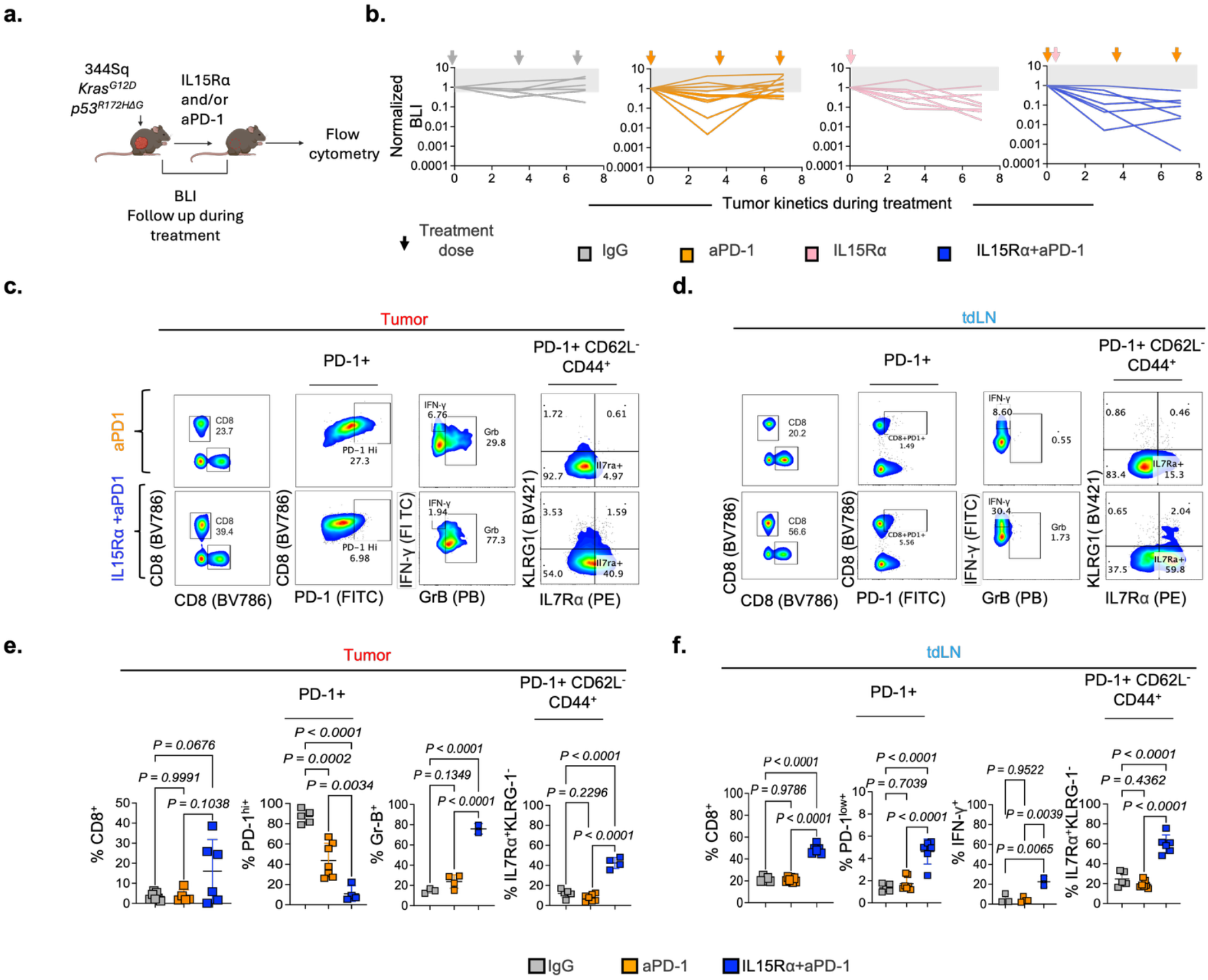
IL-15 superagonist results in robust tumor control and induces a memory-precursor effector-like T cell (Tmpe) phenotype in vivo. a. Experimental design. b. BLI plots of primary tumor burden kinetics during in vivo treatment with IgG (gray), aPD-1 (orange), IL-15 superagonist (pink) or aPD-1 + IL-15 superagonist (blue). c, d. Representative flow cytometry plots of CD8+ T cells in the tumor and tdLN of mice from the different treatment groups. e, f. Flow cytometry quantification of CD8+ T cells treated with IgG (gray), aPD-1 (orange), IL-15 superagonist (pink) or aPD-1 + IL-15 superagonist (blue) after treatment, corresponding to the plots shown in c and d. Statistics used were ANOVA (e, f).

Further characterization of CD8 T cell responses across compartments showed that, in the tdLN, IL-15 superagonist in combination with aPD-1 therapy increased the percentage of PD-1^+^ effector/effector memory T cells expressing IL-7Rα^+^(59.8% vs. 15.3%) and exhibiting cytotoxic features, demonstrated by increased IFN-γ expression (22% vs. 7%) compared with aPD-1 monotherapy. Similar phenotypic changes were observed in the primary tumor, where CD8 T cells were predominantly represented by an effector subtype. Notably, CD8^+^ T cells from combination-treated mice displayed a lower proportion of PD-1^hi^ T cells (39.5% vs. 82.3%), increased expression of granzyme B (90% vs. 40%), and maintained a higher proportion of IL-7Rα□ cells (40.9% vs. 4.97%) compared with aPD-1 alone **(Fig. 1c-f)**. Together, these findings show that IL-15 superagonism promotes a proportional expansion of a T cell subset with cytotoxic features that maintains IL-7Rα□ expression and is KLRG1^-^, gaining a Tmpe-like phenotype.

### Single cell sequencing reveals changes in CD8^+^ T cell clonal dynamics induced by IL-15 superagonism *in vivo*

To further understand CD8^+^ T cell tumor-specific responses induced by treatment, we performed paired single-cell T cell receptor (TCR) and RNA sequencing following previously published methods [37, 38]. We sorted CD45 CD3 CD8 T cells from tdLNs and tumors from mice treated with IgG, aPD-1, IL-15 superagonist, or combination therapy (**Fig. 2a**). By leveraging TCR CDR3 sequences, we identified tumor relevant expanded clones (cell number n>2) present in the tdLN by TCR match with the tumor or tumor matched (TM) clones. This approach enabled us to identify and interrogate a wide spectrum of naturally arising clones across the breadth of affinities of the antitumor response and the effect of IL-15 superagonist on CD8+ T cell clonal dynamics. TM clones constituted approximately 23% of all CD8 T cells (total n = 20,223 TM cells), indicating a substantial degree of clonal overlap between the tumor and tdLN repertoires **(Supplementary Fig. 2a)**.

**Figure 2.**
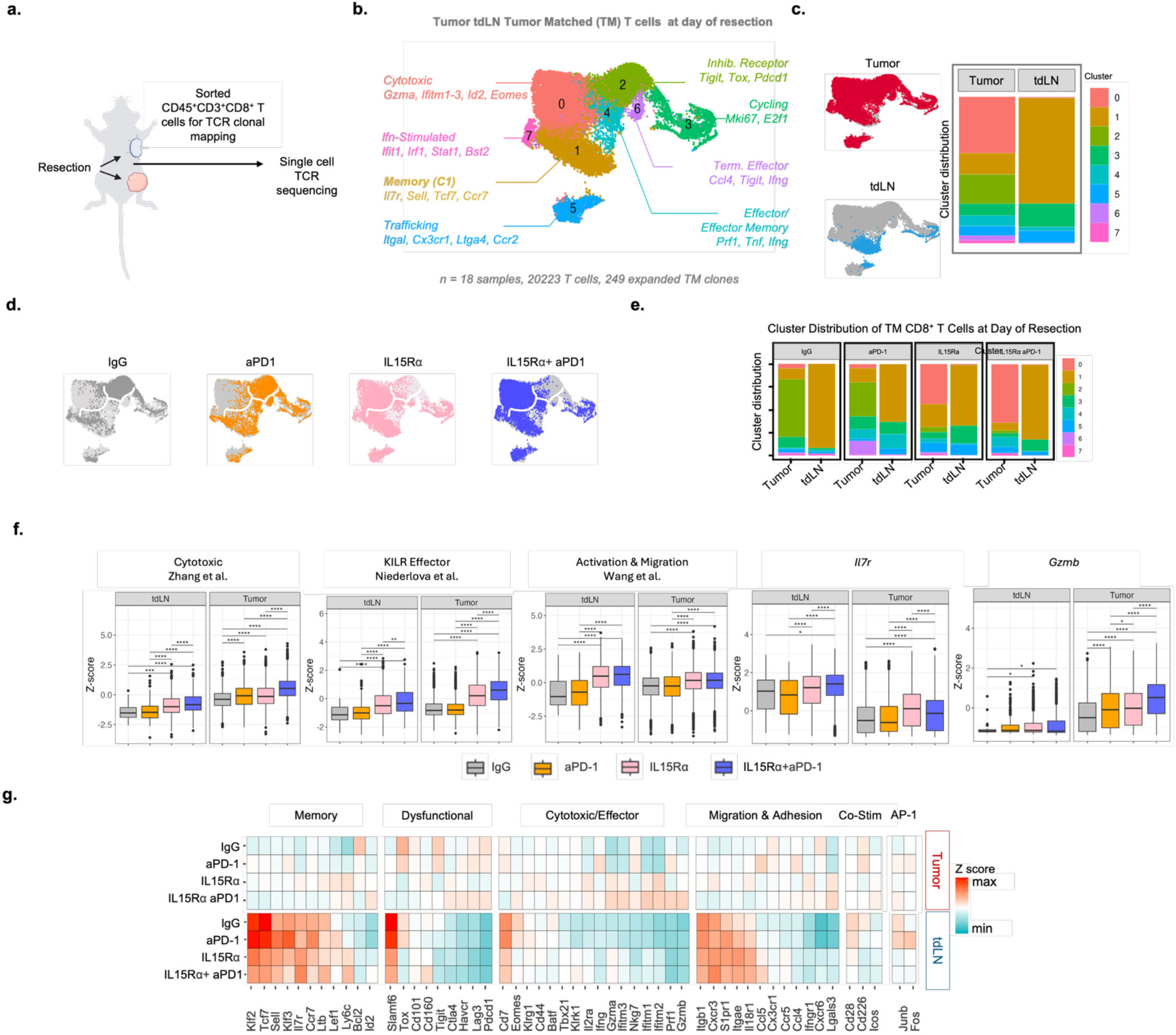
Single cell sequencing reveals changes in CD8+ T cell clonal dynamics induced by IL-15 superagonism in vivo. a. Experimental scheme. b. Clustering and UMAP visualization of CD8+ tumor-matched (TM) clones from tumor and tdLNs at the day of resection (n = 249 clones), integrated from 9 mice across two independent experiments; colors denote transcriptional clusters, labeled with functional annotations. c. UMAP showing cluster distribution of tumor-matched clones from tumor and tdLN (left), and cluster distribution for each cluster in b in the respective tissues (right). d. UMAP showing clone distribution per treatment. e. Cluster distribution for each cluster in b per treatment and tissue. f. Z score for cytotoxic, KLRK effector, activation and migration, Il7r and Gzmb signatures per treatment. g. Heatmap for genes grouped by function (memory, dysfunctional, cytotoxic/effector, migration and adhesion, co-stimulation (Co-Stim) and AP-1), with average Z score by tissue and treatment for each gene. Statistics used were t-test (f).

Unsupervised UMAP clustering of expanded TM clones identified eight transcriptionally distinct subsets spanning from the less differentiated, memory-like to the more exhausted, terminally differentiated states (**Fig. 2b**). Memory-enriched clusters were characterized by high expression of trafficking and homeostatic genes (*Sell, Ccr7, Klf2, Tcf7*) [38], and minimal expression of cytotoxic or inhibitory receptor genes. In contrast, activated and exhausted-like subsets exhibited increased expression of cytolytic mediators (*Gzmb, Gzmf, Prf1, Ifitm1, Ifitm2, Ifitm3, Nkg7, Klrk1*) [39] and inhibitory receptors (*Tox, Havcr2, Pdcd1, Entpd1, Lag3, Ctla4*) [13]. These transcriptional patterns were consistent with established T cell differentiation signatures **(Supplementary Fig. 2b)**.

TM clones exhibited marked tissue-dependent transcriptional clustering. Within the tdLN, 72% of TM clones resided within a single memory-like cluster, whereas in the tumor, clones were more widely distributed, with 47% localized within activated clusters (clusters 0, 2, 4, 6, and 7) (**Fig. 2c**). Differential expression analysis revealed that tdLN TM T cells preferentially expressed genes linked to early differentiation and memory formation (*Tcf7, Klf2, Il7r, Ly6c2*) [40]. In contrast, tumor-infiltrating clones upregulated transcripts associated with activation, cytotoxicity, and inhibitory receptor engagement (*Gzmb, Prf1, Ifng, Havcr2, Pdcd1, Ctla4*) [41]. Consistent with these transcriptional differences, modular scoring showed that tdLN TM clones displayed a naïve/central memory-like signature, whereas their tumor-infiltrating counterparts exhibited increased effector and terminal exhaustion scores (**Supplementary Fig. 2b, c**). TM clones in the tdLN expressed Tcf7 and *Il7r*, markers of long-lived memory [42], while tumor-infiltrating clones displayed high *Pdcd1* and *Havcr2* expression, indicative of progressive differentiation and exhaustion.

Examining CD8□ T cell distribution revealed significant treatment-dependent shifts. Within the primary tumor, IL-15 superagonist treatment, either alone or in combination with PD-1 blockade, enriched clones within a cytotoxic cluster (cluster 0) characterized by elevated expression of *Gzma, Id2,* and *Eomes.* In contrast, TM clones from mice treated with PD-1 blockade or IgG alone predominantly localized clones to cluster 2, which exhibited high inhibitory receptor expression (*Tigit, Tox, Lag3, Pdcd1*) [43]. Within the tdLN, all treatment regimens showed most TM clones were primarily located within the memory-like cluster 1 while also increasing representation in clusters 3 and 5, which were enriched for chemotaxis and proliferation-associated gene programs **(Fig. 2d, e)**. Notably, IL-15 superagonism and PD-1 blockade induced distinct transcriptional programs in the tdLN, suggesting divergent mechanisms of immune modulation. PD-1 blockade preferentially upregulated AP-1 family transcription factor expression in clones present in the tumor and memory-associated genes such as *Lef1, Ccr7,* and *Sell* in clones present in the lymph node. In contrast, IL-15 superagonist therapy drove a transcriptional profile marked by heightened cytotoxicity, activation, and migratory potential **(Fig. 2f, g**).

Collectively, these findings demonstrate that IL-15 superagonist therapy, alone or in combination with PD-1 blockade, promotes a highly cytotoxic/activated phenotype, whereas PD-1 blockade alone preferentially enriches for clones with high inhibitory receptor expression. Furthermore, clones within the tdLN retain a progenitor-like, memory-like associated state, serving as a reservoir for differentiated, cytotoxic effectors within the tumor, a process that is further potentiated by IL-15 superagonism.

### IL-15 superagonist increases clonal diversity of tumor-relevant T cells with a Tmpe-like phenotype at the tumor-draining lymph node level

It has been shown that increased diversity of CD8^+^ T cell clones overlapping between the tumor and the tdLN is associated with superior antitumor responses after CD4 depletion and ICB therapy [44, 45]. Additionally, it is known that IL-15 promotes T cell survival [46]; thus, we hypothesized that IL-15 superagonism would increase the diversity of tumor relevant clones. T cell clonal dynamics analysis of TM clones showed that mice receiving aPD-1 or IL-15 superagonist therapy exhibited increased TM clonal diversity relative to isotype controls (14 clones), averaging 22 and 45 TM clones, respectively, while combination therapy yielded 43 TM clones **(Fig. 3a)**. These findings indicate that while both PD-1 blockade and IL-15 superagonist broaden the CD8^+^ T cell repertoire, IL-15 notably enhances clonal diversity. Further analysis revealed that memory-enriched cluster 1 exhibited the greatest increase in clonal diversity, with 90% of tdLN-localized TM clones residing within this subset. IL-15 superagonist-treated and combination-treated mice exhibited 28 and 24 unique clones, respectively, compared with only 3 and 5 in IgG- and aPD-1-treated mice, respectively **(Fig. 3b, c)**. These data suggest that IL-15 superagonist expands clonal diversity primarily within a progenitor-like subset in the lymph node.

**Figure 3.**
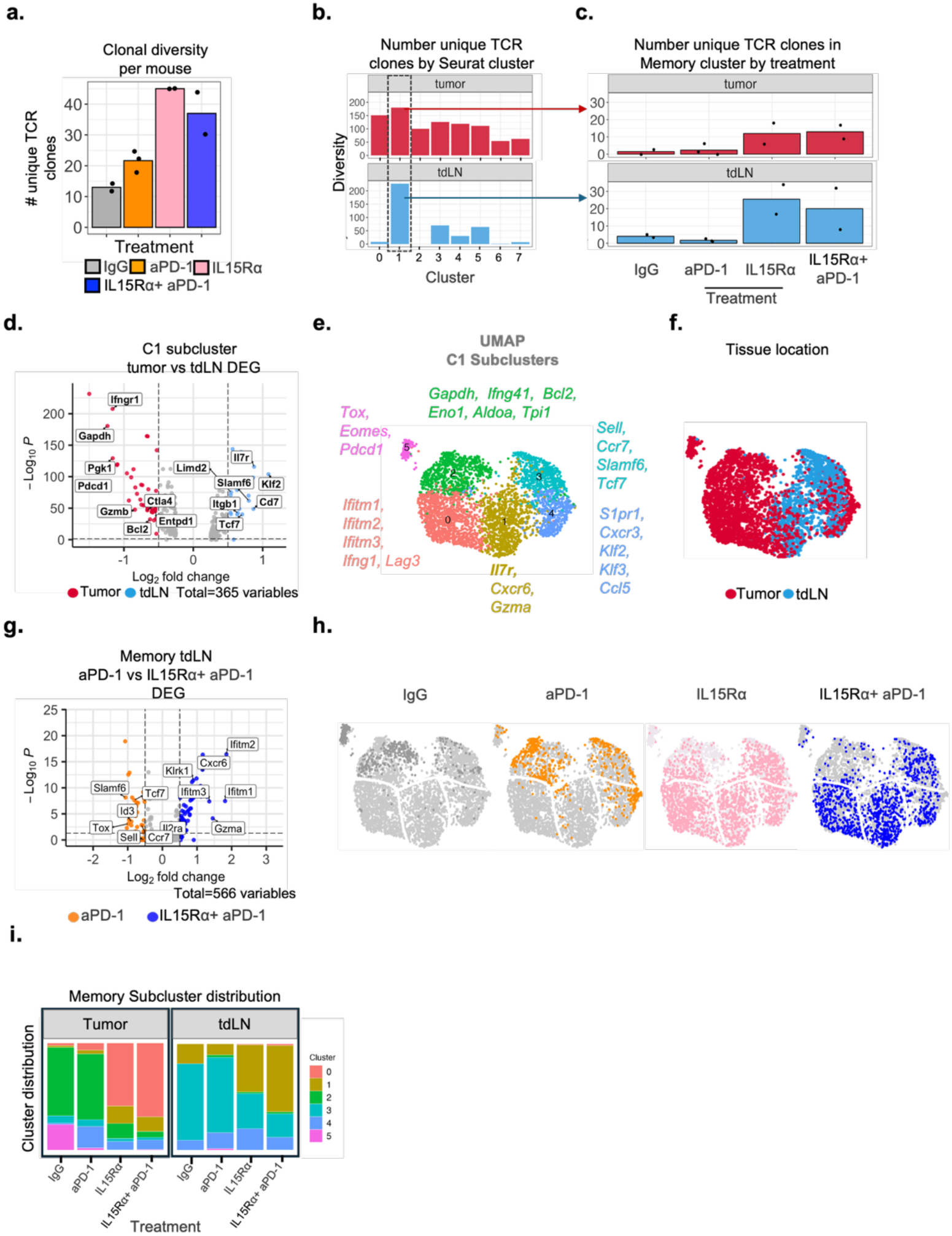
IL-15 superagonist increases clonal diversity of tumor-relevant T cells with a Tmpe-like phenotype at the tumor-draining lymph node level. a. Quantification of the number of unique TCR clones per mouse and treatment; each dot represents one mouse (n = 9). b. Bar plots showing clonal diversity in the tumor (top) and the tumor-draining lymph node (bottom) per cluster depicted in Fig. 2e. c. Number of unique TCR clones in the memory cluster (cluster 1) per treatment in the tumor (top) and tumor-draining lymph node (bottom). d. Volcano plot of differentially expressed genes between tumor and tdLN memory T cells. e. UMAP of memory subclusters derived from memory cluster 1 in Fig. 2b, labeled by the top genes within each subcluster. f. UMAP showing distribution of tumor (red) and tdLN (blue) T cell clones across memory subclusters. g. Volcano plot of differentially expressed genes of tdLN memory T cells from mice treated with aPD-1 vs. aPD-1 + IL-15 superagonist. h. Clone distribution in memory subclusters by treatment. i. Bar plots showing cluster distribution in memory subclusters by tissue and treatment.

To further dissect the heterogeneity within the memory cluster, we identified its constituent subclusters (C1 subcluster) [47-49]. Differential expression analyses revealed that tdLN clones within this memory state exhibited a progenitor-like phenotype (*Klf2, Il7r, Tcf7*), whereas tumor-infiltrating clones expressed genes associated with cytotoxicity and activation (*Gzmb, Pdcd1, Ifngr1*) (**Fig. 3d**). Memory T cells were further subdivided into five transcriptional subclusters (SC) based on the highest relative genes expressed: early cytotoxic effector/effector memory-like (SC0), *Il7r* high (SC1), glycolysis-associated (SC2), naïve/central memory/progenitor-like (SC3), and *Klf2* high trafficking/activation memory-like (SC4) (**Fig. 3e**). Most tdLN T cells were concentrated in early differentiation states (SC1, SC3, SC4), while T cell clones in the tumor concentrated in subclusters SC0, SC2, and SC4 (**Fig. 3f**). Differential expression analysis between PD-1 and combination therapy within tdLN T cells revealed that PD-1 blockade resulted in enrichment of progenitor/exhausted-like genes (*Sell, Tcf7, Tox, Slamf6*), whereas combination therapy enriched for cytolytic markers (*Ifitm1, Ifitm2, Ifitm3, Gzma, Klrk1*) (**Fig. 3g**), with IL-15 superagonist treatment preferentially increasing the proportion of SC1 T cell clones (**Fig. 3h, i**). By contrast, PD-1-treated and IgG-treated memory T cells remained predominantly in SC3 and exhibited a stem-like signature [50].

These transcriptional differences extended to tumor-infiltrating T cells, where IL-15 superagonist therapy increased the representation of SC0 relative to non-IL-15 superagonist-treated mice. These data suggest that IL-15 superagonism induces an effector phenotype with cytotoxic features and memory-precursor-like features that is enriched in the tdLN, and may shape tumor effector function upon tumor infiltration.

### IL-15 superagonist Tmpe-like phenotype is dependent on CD122 receptor subunit and induces robust STAT5 phosphorylation

To explore the underlying signaling responsible for the robust CD8^+^ T cell phenotype we observed in vivo, we established an in vitro system using activated OT-1 CD8^+^ T cells **(Fig. 4a).** T cells were treated with IL-15 superagonist, aPD-1, a combination of IL-15 superagonist/aPD-1, or IgG. Flow cytometry phenotyping at day 3 of culture showed that CD8^+^ T cells treated with IL-15 superagonist maintained a relatively lower proportion of PD-1^+^ T cells compared with aPD-1- or IgG-treated groups **(Supplementary Fig. 3a-c)**.

**Figure 4.**
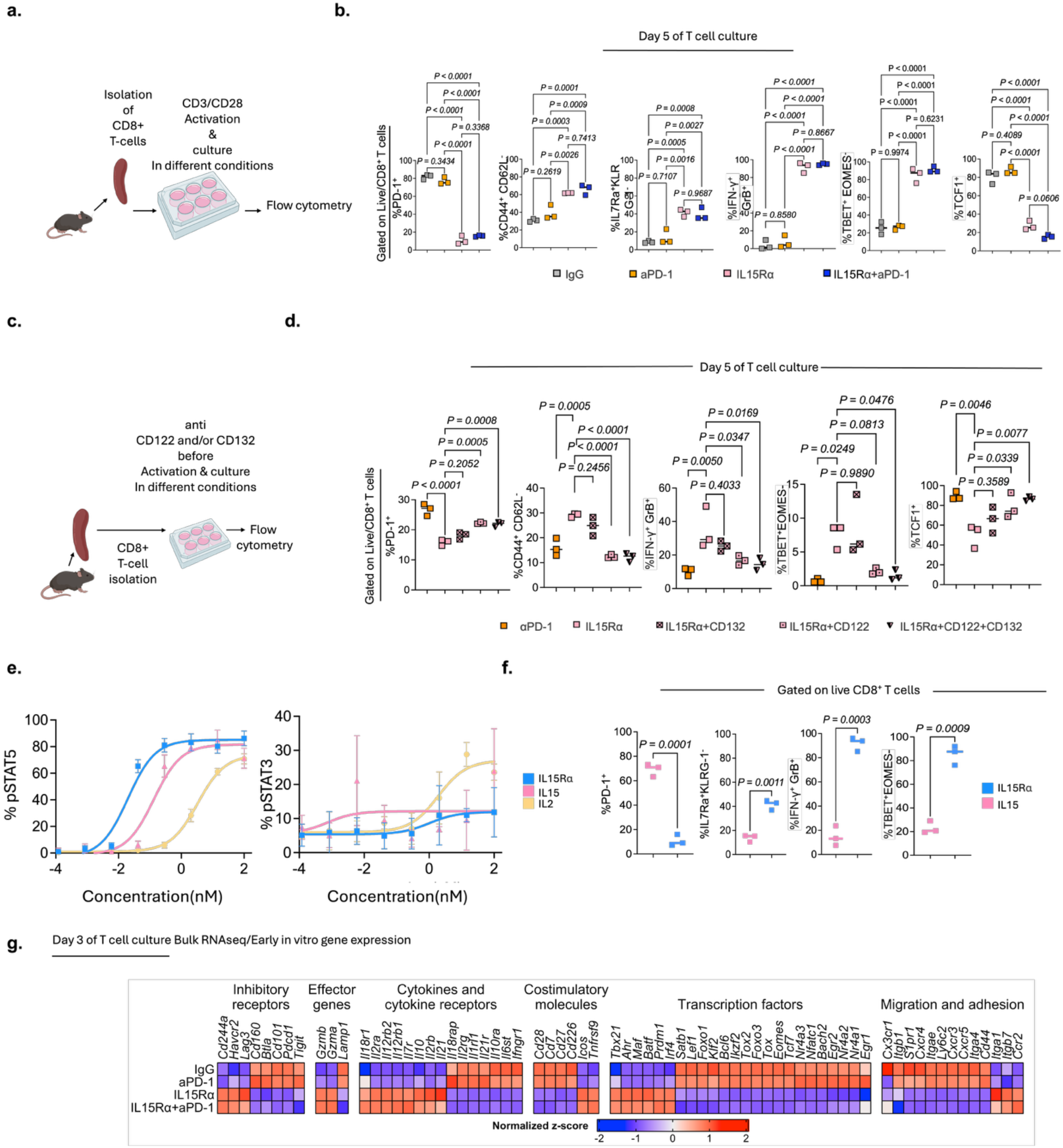
IL-15 superagonist Tmpe-like phenotype is dependent on CD122 receptor subunit and induces robust STAT5 phosphorylation. a. Experimental design. b. Flow cytometry quantification of CD8+ T cells treated with IgG (gray), aPD-1 (orange), IL-15 superagonist (pink) or aPD-1 + IL-15 superagonist (blue) at day 5 of culture. c. Experimental design. d. Flow cytometry quantification of CD8+ T cells treated with aPD-1 (orange) or IL-15 superagonist (pink) at day 5 of culture after CD122 and/or CD132 blockade. e. IL-15 superagonist, IL-15 and IL-2 dose-response curves showing the percentage of mouse CD8+ T cells that are pSTAT5+ (left) or pSTAT3+ (right). Error bars indicate mean ± SD of triplicate wells. f. Flow cytometry quantification of IL-15 superagonist and IL-15 treated CD8+ T cells at day 5 of T cell culture. g. Heatmap for genes grouped by function (inhibitory receptor, effector genes, cytokines and cytokine receptors, costimulatory molecules, transcription factors, migration and adhesion), with average Z score by treatment for each gene. Statistics used were one-way ANOVA (b, d) and t-test (f).

By day 5, IL-15 superagonist significantly increased the proportion of CD8^+^ T cells that were CD62L^-^CD44^+^ and displayed a Tmpe-like phenotype, defined by IL-7Rα^+^/KLRG1^-^ expression. IFNγ and GzmB expression were also significantly higher, when compared with aPD-1 treatment and untreated controls, while maintaining a lower proportion of PD-1^+^ CD8^+^ T cells relative to other treatment groups **(Fig. 4b)**, recapitulating the Tmpe-like phenotype observed in vivo.

Clinical trials using different modified cytokines have shown that these novel molecules can engage with their receptor differently from their natural counterparts; an example of this is genetically modified IL-2 molecules currently under clinical testing that do not bind to CD25, in contrast with canonical IL-2 [19, 51]. Thus, we interrogated which receptor subunits were required for IL-15 superagonist-mediated changes in CD8^+^ T cell phenotype.

Canonical IL-15 signaling occurs through engagement of the heterotrimeric receptor composed of IL15Rα (CD215), IL2Rβ (CD122), and IL2Rγ (CD132), leading to downstream JAK activation and STAT5 phosphorylation (pSTAT5) [21]. To investigate which receptor subunits were involved in the observed phenotype, we inhibited CD122, CD132, or both before treatment with the IL-15 superagonist, as this immunocytokine already incorporates the IL-15Rα (CD215) chain to the IL-15 molecule. Flow cytometry analysis at day 5 of T cell culture revealed that CD122 blockade was sufficient to abrogate IL-15 superagonist-induced effector phenotype (**Fig. 4c, d**), suggesting that the effect of the IL-15 superagonist on CD8^+^ T cells is mostly dependent on CD122.

Because engineered or surrogate cytokines can bias downstream STAT activation [52, 53], we evaluated whether STAT phosphorylation differed between IL-15 superagonist, standard IL-15, and IL-2. Phosphorylation flow cytometry experiments demonstrated that IL-15 superagonist induced a greater magnitude of pSTAT5 signaling compared with conventional IL-15 and IL-2, and induced minimal detectable STAT3 phosphorylation, unlike IL-2 (**Fig. 4e**). We hypothesized that these differences in downstream signaling were associated with differences in phenotype compared with canonical IL-15. Flow cytometry on day 5 of culture of cells treated with canonical IL-15 or IL-15 superagonist showed significant differences in T cell phenotype (**Fig. 4f**), suggesting that downstream bias in STAT signaling may explain these phenotypic differences.

pSTAT5 signaling promotes the expression of effector function-related genes [54], while also playing a critical role in memory formation [55, 56] and subverting “exhaustion” by antagonizing NFAT transcription factor activity on PD-1 promoter [57]. Consistent with this, early gene expression changes evaluated by bulk RNAseq at day 3 after in vitro treatment with IL-15 superagonist showed relatively lower expression of *Pdcd1*, and increased expression of genes such as *Il7r*, *Il2ra*, *Il2rb*, and effector genes such as granzyme B and interferon γ [55, 58] (**Fig. 4g**). This was consistent with phenotypic differences observed at days 3 and 5 of culture, and suggests that the generation of a Tmpe-like phenotype induced by this IL-15 superagonist might be the result of increased STAT5 phosphorylation and signal transduction.

### IL-15 superagonist results in lower metastatic recurrence and promotes systemic persistence of tumor-relevant memory CD8^+^ T cell clones at non-draining lymph nodes

Our highly metastatic model gave us the opportunity to evaluate differences in metastatic recurrence and memory induced by treatment. Mice received aPD-1, IL-15 superagonist, combination therapy, or IgG. Tumors were resected by survival surgery 3 days after the last dose of aPD-1, and mice were followed up by bioluminescence imaging for metastasis. Longitudinal imaging over 200 days revealed that IL-15 superagonist in combination with PD-1 blockade substantially diminished metastatic recurrence and improved overall survival (**Fig. 5a**), with around 75% of mice remaining alive at day 200, in contrast with the IL-15 superagonist or aPD-1 single-treatment groups, where only 14% and 18% of the mice survived, respectively (**Fig. 5b**).

**Figure 5.**
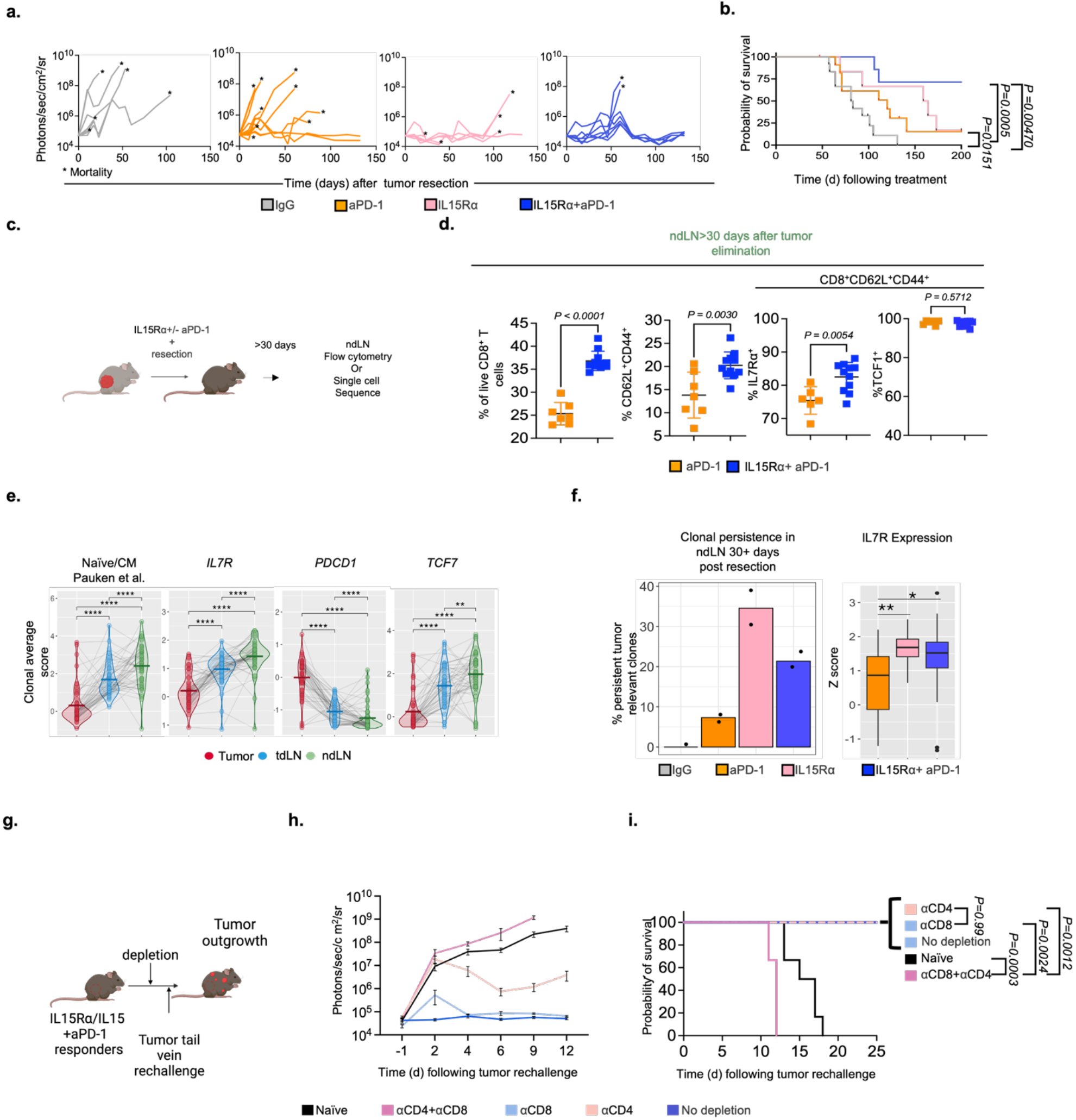
IL-15 superagonist results in lower metastatic recurrence and promotes systemic persistence of tumor-relevant memory CD8+ T cell clones at the non-draining lymph nodes. a. BLI plots of metastatic disease in mice from the different treatment groups after primary tumor resection, up to 200 days. Each dot represents a mouse mortality event. b. Kaplan Meier survival plots. c. Experimental design. d. Flow cytometry quantification of CD8+ T cells from the inguinal non-draining lymph node, resected more than 30 days after primary tumor resection, from mice under neoadjuvant aPD-1 or combination therapy, showing percentages of central memory (CD62L+CD44+) CD8+ T cells. e. Average Z score for naïve/CM (Pauken et al.), Il7r, Pdcd1 and Tcf7 expression per clone; each line traces and connects matching clones in the tumor (red), tdLN (blue) and 30+ day ntdLN (green). f. Percentage of persistent tumor-relevant clones in the ntdLN more than 30 days after resection of the primary tumor, identified by TCR match. g. Schematic design of the depletion and tumor rechallenge experiments in combination-treated long-term responders. h. BLI plots of tumor burden in the chest after tail-vein injection of combination-treated, recurrence-free mice with no depletion (blue) or receiving different depleting antibodies: CD4 (yellow), CD8 (purple), CD4/CD8 (green), and naïve age-matched mice (black). i. Overall survival for the experimental groups in h. Statistics used were t-test (d, e, f) and Šidák-corrected log-rank (Mantel Cox) test (b, i).

As IL-15 has a role in memory formation and T cell survival, we hypothesized that mice treated with IL-15 superagonist would have an increase in central memory T cells (TCM). To investigate whether there were differences in memory formation among long-term survivors from different treatments, we resected the contralateral ntdLN (inguinal or axillary) of tumor-free mice after 30 and up to 120 days post-treatment for evaluation of systemic memory **(Fig. 5c)**.

Flow cytometry immunophenotyping of ntdLNs at this timepoint showed that mice that received IL-15 superagonist in combination with aPD-1 therapy still had a significant proportional expansion of CD8 T cells when compared with aPD-1 (37% vs. 25%), a change observed initially after treatment, suggesting persistent changes in the CD8^+^ T cell compartment. The presence of a memory population defined by the expression of CD62L^+^, CD44^+^ and TCF-1^+^ was observed regardless of the treatment group and suggested that these mice had entered a memory state after tumor exposure/eradication. However, mice that received IL-15 superagonist in combination with aPD-1 therapy showed a significant increase in TCM cells (20% vs. 14%) compared with PD-1 blockade, and markedly enhanced the proportion of IL-7Rα^+^ T cells within the TCM subset (84% vs. 76%) **(Fig. 5d, Supplementary Fig. 4a, b)**.

To determine whether this expansion in memory reflected an increase in persistence of tumor-relevant T cells, we resected the ntdLN (axillary or inguinal) more than 30 days after primary tumor resection from mice whose tumors and tdLNs had been previously sequenced at initial time of treatment. We conducted scTCR and RNA sequencing so that we could identify tumor relevant clones present in the tumor and match to persisting clones in the ntdLN by TCR sequence.

Analysis at the level of individual persistent clones revealed similarities between ntdLN and tdLN (Supplementary Fig. 4a, c), while tumor clones exhibited decreased expression of memory-associated genes **(Fig. 5e)**. Clonal-average scores showed that tumor-infiltrating clones had relatively reduced quiescence and memory signatures compared with ntdLN and tdLN counterparts **(Supplementary Fig. 4d-f)**.

Clonotypic tracking by treatment groups demonstrated that neoadjuvant therapy markedly enhanced the persistence of tumor-matched clones in ntdLNs beyond day 30, with IL-15 superagonist alone or in combination with aPD-1 doubling the proportion of persistent clones compared with aPD-1 monotherapy. In the IgG control group, almost no persistent clones were found **(Fig. 5f)**. Importantly, clones in IL-15 superagonist and combination therapy groups displayed heightened expression of IL-7Rα compared with PD-1 blockade alone (**Supplementary Fig. 4g)**.

The development of long-term protective CD8^+^ T cell immunity has been linked with lower metastatic relapse [59]. As we observed increased persistence of tumor-relevant CD8+ T cell clones in IL-15 superagonist treated mice, we hypothesized that memory protection induced by this superagonist was primarily dependent on CD8^+^ T cells. To explore this, we implanted mice with 344SQ tumor cells in the right flank and subsequently treated them with IL-15 superagonist in combination with aPD-1 as previously described. Residual tumor was resected through survival surgery after treatment, and mice were followed up for development of metastatic disease. Thirty days after primary tumor resection, metastasis-free mice underwent depletion of different immune subsets and were subsequently rechallenged with the same tumor cell line by tail vein injection (**Fig. 5g**). Surprisingly, CD8^+^ depletion was not sufficient to abrogate protection during rechallenge, and only combined depletion of CD4^+^ and CD8^+^ T cells resulted in loss of protection at the memory stage (**Fig. 5h, i**). This “redundant” memory was not observed in aPD-1 treated mice, where both subsets were simultaneously required for protection during rechallenge (**Supplementary Fig. 4h**).

To explore whether IL-15 superagonist also induced early changes in the CD4^+^ compartment, we conducted flow cytometry of CD4^+^ T cells at the time of treatment. Immune phenotyping demonstrated that IL-15 superagonist also induced phenotypic changes in this T cell subset, significantly increasing the expression of effector molecules such as granzyme B and interferon γ at the tumor and tdLN respectively (**Supplementary Fig. 4g**), suggesting enhanced functionality in these populations as well as potential for cytotoxic activity. In summary, these findings indicate that the initial response to IL-15 superagonist also engages CD4^+^ T cells and results in a lasting unique memory where either CD4^+^ or CD8^+^ T cells alone can mediate memory protection.

## Discussion

Using a clinically relevant murine model of lung cancer with metastatic relapse, we provide a framework to explore the effects of an IL-15 superagonist on antitumor immunity. We found that IL-15 superagonist alone or in combination with aPD-1 results in increased primary tumor control and long-lasting anti-tumor immunity.

IL-15 superagonist induced a proportional expansion of CD8^+^ T cells at the tumor and tdLN level, and increased the proportion of T cells with a Tmpe-like phenotype, characterized by CD62L^-^ CD44^+^ IL-7Rα^+^ KLRG1^-^ expression and increased cytokine expression (IFN-γ, granzyme B). This phenotype was dependent on the tissue localization, with tumor-draining lymph node CD8^+^ T cells retaining a more functional phenotype after therapy and in contrast to matching clones present in the tumor, suggesting that IL-15 superagonist may rely on precursor CD8^+^ T cells from the tdLN for an optimal anti-tumor response.

Harnessing cytokines to boost antitumor responses has regained recent interest, especially with novel engineering strategies that have decreased toxicity and target the tumor microenvironment (TME) [60, 61]. Furthermore, there is ongoing exploration of STAT signaling downstream of different types of cytokine receptors to reprogram immune responses [62]. IL-15 superagonist required engagement of the IL2Rβ chain subunit (CD122) and resulted in robust phosphorylation of STAT5, with minimal STAT3 engagement, in contrast with standard IL-15 or IL-2, with changes at the RNA level that included relatively lower expression of *pdcd1* and increased relative expression of *il7r* and cytokine genes supporting the phenotype observed at different timepoints during in vitro and in vivo experiments. All this work underscores the importance of cytokine signaling rewiring to modulate T cell differentiation and exhaustion to generate more functional and therapeutically suitable T cell phenotypes, and, alongside other authors, supports a new therapeutic avenue focused on STAT signal manipulation to induce different T cell states that can display variable levels of fitness in different tumor contexts [62].

Importantly, we also explored how modified cytokines can induce long-lasting T cell responses and considerably increase the persistence of tumor-relevant clones at the memory stage. IL-15 superagonism generated a distinct form of memory. Surprisingly, during tumor rechallenge, single depletion of either CD8 or CD4^+^ T cells failed to abrogate protection, whereas combined depletion was the only condition under which mice lost the ability to reject the tumor rechallenge. Addtionally, in aPD-1 treated mice, single depletion of either subset was sufficient to impair protection. This redundant memory, induced by IL-15 superagonist in combination with aPD-1, was associated with lower metastatic recurrence. How these changes in memory response occur, what are the specific determinants for persistence of these tumor-relevant clones, and how these processes contribute to reduced metastatic reccurence remain to be understood.

IL-15 superagonist immunomodulation has emerged as a safe and potent strategy to enhance endogenous anti-tumor immunity. The phase I clinical trial of KD033 in patients with metastatic and locally advanced disease (NCT04242147) reported no dose-limiting toxicities [63]. The IL-15 superagonist N-803, now FDA-approved for BCG-unresponsive early-stage bladder cancer, achieved a complete response in 71% of patients [64], and is currently being tested in patients with advanced NSCLC (NCT05096663). Here, we show that IL-15 superagonist alone or in combination with PD-1 blockade elicits potent tumor-directed cytolytic activity and sustains long-lived, tumor-specific T cells. These findings provide a strong rationale for cytokine-based therapies such as this superagonist to achieve durable clinical benefit in solid tumors.

## Limitations

This study has limitations. The in vivo work was performed in a single syngeneic model of lung adenocarcinoma, chosen because it is resectable and spontaneously metastatic and therefore permits clonal tracking from treatment through surgery to relapse. Althought Martomo et al. have shown that IL-15 superagonism results in significant primary tumor control in different types of tumors, further T cell phenotype characterization and memory persistence needs to be explored in other cancer types. Cytotoxic function was inferred from *in vivo* primary tumor control, granzyme B and interferon γ expression and from transcriptional cytotoxicity scores rather than measured directly, so the killing capacity of the Tmpe population remains to be established. The association between STAT5 phosphorylation and phenotype was defined in vitro and is correlative, since CD122 blockade establishes receptor dependence but STAT5 itself was not genetically manipulated. Tumor relevance of clones was assigned operationally by TCR sharing between tumor and lymph node rather than by antigen specificity testing. Finally, the basis of the redundant CD4 and CD8 memory, including whether CD4 T cells act through acquired cytotoxicity and/or through help, was not determined.

## List of Abbreviations

ACK: ammonium-chloride-potassium
ADCC: antibody-dependent cellular cytotoxicity
ANOVA: analysis of variance
aPD-1: anti-programmed cell death protein 1
BLI: bioluminescence imaging
CDR3: complementarity-determining region 3
DAPI: 4’,6-diamidino-2-phenylindole
FACS: fluorescence-activated cell sorting
GzmB: granzyme B
IACUC: Institutional Animal Care and Use Committee
ICB: immune checkpoint blockade
IFN-γ: interferon gamma
IL: interleukin
IRB: Institutional Review Board
JAK: Janus kinase
KILR: KLRK1 IL-7Rα effector
KLRG1: killer cell lectin-like receptor G1
NSCLC: non-small cell lung cancer
ntdLN: non-tumor-draining lymph node
PD-1: programmed cell death protein 1
PD-L1: programmed death ligand 1
PMA: phorbol 12-myristate 13-acetate
pSTAT: phosphorylated signal transducer and activator of transcription
SC: subcluster
scRNA-seq: single-cell RNA sequencing
scTCR-seq: single-cell T cell receptor sequencing
STAT: signal transducer and activator of transcription
TCM: central memory T cell
TCR: T cell receptor
tdLN: tumor-draining lymph node
TM: tumor-matched
TME: tumor microenvironment
Tmpe: memory-precursor effector-like T cell
Tpex: progenitor exhausted T cell
UMAP: uniform manifold approximation and projection.

## Consent for publication

Not applicable.

## Availability of data and material

Single-cell RNA sequencing and single-cell T cell receptor sequencing data generated in this study have been deposited in the Gene Expression Omnibus under accession [GSE######] and will be publicly released upon publication. All other data supporting the findings of this study are available from the corresponding author upon reasonable request

## Acknowledgements

We thanks J. McCormick, T. Baumgartner and P. Byrne of the Flow Cytometry Core Facility, and Dr. Jenny Xiang of the Genomics Resources Core Facility for professional advice.

## Availability of data and material

The datasets generated and analyzed during the current study are available from the corresponding author upon reasonable request and will be deposited in the Gene Expression Omnibus prior to publication.

## Authors’ contributions

J.V.V. conceived and supervised the study. T.D.C. and J.V.V. designed the experiments. T.D.C. performed experiments, analyzed data, generated experimental figures. R.H., L.Y. and S.C. performed the computational analyses. S.A.M. provided KD033 and contributed to study design. G.J.M., B.S., M.S.T., S.E.C., G.E.R., C.G., Y.H., Y.B., J.C.R., J.P., D.O.O., M.M.M., A.S., T.E.M. contributed to experiments. O.E., N.K.A., V.M., J.C.R. and G.J.M provided supervision and suggestions. T.D.C., R.H. and J.V.V. wrote the manuscript with input from all authors. All authors read and approved the final manuscript. J.V.V. is the guarantor.

## Materials and methods

### Animal work

Animal work was performed in accordance with an animal protocol approved by the institutional Animal Care and Use Committee at Weill Cornell Medical College (protocol no. 0806-762A). Female 129S1/SvImJ mice were purchased from the Jackson Laboratory (Cat. No. 002448).

### Culturing of cell lines

344SQ lung cancer cell line was kindly provided by Dr. Jonathan Kurie’s lab (MD Anderson Cancer Center, Houston, TX). Tumor cells were obtained from subcutaneous tumors from *p53^R172H^*^Δ*g/*+^ *K-ras^LA1/^*^+^ mice. These cells were cultured in DMEM supplemented with 10% FBS, 10 mM L-glutamine and 100 U ml^−1^ penicillin with 100□μg□ml^−1^ streptomycin. mCherry-luciferase stable expression was achieved using lentiviral transduction.

### Flank injection of tumor cells in syngeneic mice (129S1/SvImJ)

We used immune-competent 129Sv (Jackson Laboratory, Bar Harbor, ME, USA 04609) females that were 8-12 weeks old for our *in vivo* studies. 1 × 10^6^ 344SQ mCherry-luciferase-expressing cells were harvested at 50-70% confluence and resuspended in 100 μl of PBS and injected in the right flank. Tumor growth in vivo was evaluated at day 3 after injection and once weekly via BLI using a Xenogen IVIS system. To cohort mice for the different treatments, BLI data were collected 1 day before treatment and mice were grouped such that similar mean tumor burden was present in each group. All treatment groups started therapy after 3 weeks of tumor growth on the same day. For treatment with aPD-1 or IgG2a mice were injected intraperitoneally with 250 μg of aPD-1 or IgG2a as previously reported [65] on days 1, 4, and 7 of treatment. Mice treated with IL-15 superagonist (KD033) received a single dose by tail vein injection (3 mg/kg) at the same time as the first dose of aPD-1. Mice that received combination therapy received a single dose of IL-15 superagonist on day 1 of treatment together with aPD-1, followed by two additional doses of aPD-1 on days 4 and 7. Three days after the last dose of treatment, all mice underwent survival surgery at the same time.

For depletion studies, matched cohorts of mice that underwent different neoadjuvant treatments and completed at least 30 days after survival surgery received CD4-, CD8-, and/or NK-depleting antibodies depending on the depletion group with the following doses: 400 μg anti-CD4 (clone GK1.5, BioXCell), 400 μg anti-CD8a (clone 2.43, BioXCell), and 50 μl of anti-asialo GM1 antibody. Respective controls were also used depending on the depletion group compensating for antibody load in respective groups. For rechallenge experiments, 700,000 344SQ mCherry-expressing cells were injected through tail vein injection. Efficacy of depleting antibodies was confirmed by flow cytometry of blood from jaw bleeds.

### Survival surgery

The animals were anesthetized utilizing isoflurane. An injection of buprenorphine extended release (Ethiqa XR) SC was given for pre-emptive analgesia immediately after the animals were anesthetized. Depth of anesthesia was monitored at least every 15 minutes throughout the procedure. Following confirmation that a suitable anesthetic plane was attained, sterile eye lubricant was applied to both eyes to prevent corneal drying during surgery. Alternating scrubs of 70% ethanol and chlorhexidine were applied three times to the area of the surgical incision. Animals were covered with a sterile drape with a hole for access to the incision and kept warm using a heating pad. Fresh sharp scissors were used to make an incision size of up to twice the diameter of the tumor through the skin directly overlaying the mass. Sterile hemostats were used to bluntly dissect the tumor and/or tdLN (inguinal) away from the underlying normal tissue. During blunt dissection, blood vessels feeding or draining the tumor were ligated. Once the tumor was separated from the surrounding tissue, it was carefully removed using sterile forceps for grasping the mass and sterile scissors to facilitate removal. Skin edges were apposed and closed with sterile wound clips. Mice were monitored during recovery from anesthesia and were followed up at the 24-, 48-, and 72-hour timepoints for signs of post-surgical complications or pain and treated according to protocol. Surgical clips were removed between day 7 and day 14.

### Mouse tissue harvest

At specific timepoints tissues were collected during survival surgery, kept on supplemented culture media at 4 °C, then tumors were ground through a 140-μm wire mesh (Cell Screen/100 mesh, Bellco Glass) then filtered through 70-μm filters. Lymph nodes were dissected and filtered through 70-μm filters. Red blood cell lysis into single cell suspension was achieved by using ACK lysing buffer and cell pellets were washed and resuspended in FACS buffer.

### Flow cytometry staining

For surface staining, samples were incubated with a fixable live/dead viability dye (Zombie Aqua, BioLegend), stained with primary antibodies in the presence of Fc blockers, washed with FACS buffer, fixed with 1% formaldehyde, washed with FACS buffer and resuspended in FACS buffer. Transcription factor staining: Samples were incubated with Zombie Aqua fixable live/dead dye, stained with primary antibodies in the presence of Fc blockers, washed with FACS buffer, fixed with Fixation/Permeabilization Buffer (eBioscience), washed with Permeabilization Buffer (eBioscience), stained with primary antibodies for intracellular markers in Permeabilization Buffer, washed with Permeabilization Buffer and resuspended in FACS buffer.

Cytokine staining: Single cell suspension obtained after ACK lysis buffer was incubated at 37□°C in a humidified incubator for 5 hours using the Golgi blockers brefeldin A (BioLegend) and monensin (BioLegend), and cytokine stimulants PMA and Ionomycin. After incubation, cells were stained for surface and intracellular markers as described above.

pSTAT signaling assays: CD8 T cells were isolated from the spleen of OT-1 or wild-type C57BL/6 mice and cultured on CD3/CD28-coated plates in mouse T cell media supplemented with 100 U/ml IL-2 for 48 hours. After this, T cells were transferred to non-coated plates and resuspended on T cell media supplemented with IL-2 for another 48 hours. Upon completion of this time, T cells were washed and resuspended on T cell media with no cytokines for 18 to 24 hours before pSTAT signaling assays. Before stimulation, T cells were stained for Zombie and with fluorescent antibodies targeting CD3 and CD8 surface markers. Ligands were added to cells at indicated concentrations and incubated in serum-free media at 37°C for 15 to 20 min, followed by fixation with 1.5% paraformaldehyde. Cells were washed and then permeabilized with ice-cold methanol at −80°C. The cells were then washed and stained with fluorescent pSTAT detection antibodies for 2 hours, followed by analysis via flow cytometry [62].

### Sorting of T cells for single cell TCR sequencing analysis

At specific timepoints tissues were collected during survival surgery, kept on supplemented culture media at 4 °C, tissues were collected, processed and ACK lysed as described above, single cell suspension was stained using primary antibodies in the presence of Fc blockers, washed and resuspended in FACS buffer and stained with CD3, CD8, CD45, CD62L and CD45RA (human samples) or CD44 (mouse experiments) and DAPI 0.2 μg ml-1. The sorting strategy used was CD45+, CD3+, CD8+, CD62L+, and CD45RA/CD44 was used to verify phenotypes during sorting.

### scRNASeq and scTCRSeq demultiplexing and read processing

To align raw single cell RNA sequencing and single cell V(D)J sequencing data we used Cell Ranger version 3.0.2 multi. The GEX mm10-2020-A mouse reference genome was used as the transcriptional reference, along with the GRCm38 mouse reference for V(D)J. Aligned transcriptional data were processed with Read10X in R version 3.6.1, and Seurat version 3 was used to create a Seurat object with min.cells set to 3 and min.features set to 200, excluding genes detected in fewer than three cells and cells with fewer than 200 detected features. TCR V(D)J data were then processed by combining the filtered_contig_annotations.csv and clonotypes.csv datasets to recover the paired CDR3 alpha and beta amino acid sequences for each cell, which were added as metadata to the Seurat object.

### Single-cell TCR sequencing quality control and clonal analysis

CD8 T cells were retained only when at least one α and one β chain were recovered, since clonal identity cannot be assigned from a single chain. Distinct TCR clones were defined by the exact combination of CDR3 α and β amino acid sequences, and clone size was the number of cells sharing that combination. Only expanded clones, defined as clones of at least three cells, were carried into downstream clonal analysis, because clones of one or two cells cannot be distinguished from sampling noise at this sequencing depth. Clones whose cells were recovered in both the tdLN and the tumor were designated tissue-matched (tm) clones and used as the operational definition of tumor-relevant clones, on the reasoning that a TCR recovered on both sides of the tdLN to tumor axis has been engaged by antigen present in the tumor. Persisting clones, a subcategory of tm clones, were tm clones whose cells were additionally recovered in the ntdLN after tumor resection, and therefore after antigen withdrawal. Each T cell was labeled with its clone ID and tm status.

After unexpanded clones were removed, clonality within each tissue and treatment was analyzed in three ways: clonal diversity, the percentage of total tm T cells, and a normalized weighted expansion score. Clonal diversity was the total number of distinct clones within a group. The proportion of tm T cells within a given tissue and treatment was compared against the IgG-treated group for that tissue using a chi-squared test.

Clonal expansion corresponds to the size of a given clone, so comparing expansion between groups that differ in cell number requires a score that both weights larger clones more heavily and is placed on a common scale. A vector *v* of all distinct expanded clone sizes across all mice was assembled, and a linear weighting scale was defined on it such that the largest distinct clone size in *v* receives a weight of 1 and a clone size of 0 receives a weight of 0, giving the scaling constant *m* = 1/max(*v*). T cells were then partitioned into subgroups *X_j_* defined by each combination of treatment and tissue, and clone sizes were recalculated within each subgroup, so that the score reflects how expanded a tm clone is within that particular tissue rather than across the dataset as a whole. For each subgroup, the distinct clone sizes were weighted by *m*, summed, and normalized by the number of unique clones *N*, as shown in Equation 1. Weighting in this way gives heavier influence to larger, more expanded clones, so that a group dominated by a few highly expanded clones is distinguished from one containing many small clones even when the two comprise similar cell numbers.

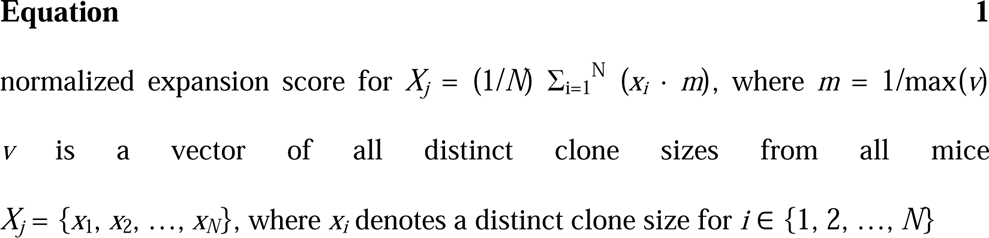

### Single-cell RNA computational processing

Single-cell RNA sequencing data were preprocessed in Seurat following the analytical framework established for paired scRNA/scTCR profiling of tumor-specific CD8 T cells [Pauken et al., Pai et al.]. Quality control removed cells whose mitochondrial gene percentage exceeded two standard deviations above the dataset mean, a data-driven threshold that excludes dying cells without imposing a fixed cutoff across samples of differing sequencing depth. Transcriptional data were normalized with NormalizeData using the LogNormalize method and a scale factor of 10,000, the 2,000 most variable features were selected with FindVariableFeatures using the vst method so that dimensionality reduction was driven by genes carrying biological variance rather than by technical depth, and principal components were computed with RunPCA. The number of dimensions carried into RunUMAP was selected from the inflection of the PCA variance plot, retaining the components that captured population structure while excluding those contributing predominantly noise.

Because every downstream analysis traces clones between tissues, inclusion in the final object required a recovered TCR, and removing cells without one served as the primary filter against non-T cells. CD8 identity in the retained cells was confirmed by expression of *Cd3d*, *Cd3e*, *Cd3g*, *Cd8a*, *Cd8b1* and *Cd8b2*; residual epithelial cells, B cells and CD4^+^CD8^−^ T cells were excluded, although after TCR-based filtering few such cells remained. Metadata for treatment group, response and recurrence status of each mouse were attached to the object and used as the grouping variables for all differential expression comparisons.

Two cohorts of mice were sequenced. Cells were integrated with Harmony on mouse ID to remove mouse-specific and sequencing-specific bias. Integration across tissues was deliberately not performed, because tumor, tdLN and ntdLN T cells are expected to differ transcriptionally as a function of tissue environment and antigen exposure, and integrating across tissue would have removed the biological signal under study. Where experiments were compared directly, a new Seurat object containing all samples was created and clustered using the same preprocessing and clustering procedure. T cells from sequenced mice had the ntdLN resected 30 days after initial resection to trace TCR clones longitudinally; mice that did not recur or die by 30 days after primary tumor resection were selected, persisting clones were identified by tracing which tm clones were also recovered in the ntdLN, and a new UMAP was generated from those persisting clones.

### Transcriptional signature analysis with TCR clonal matching category

To compare T cell functionality between clonal matching groups, relative functional scores were generated with Seurat’s AddModuleScore and then centered and scaled. Marker sets were taken from the literature: *Sell*, *Tcf7*, *Ccr7*, *Il7r*, *S1pr1* and *Klf2* as naïve and central memory markers, and *Klrg1*, *Cx3cr1*, *S1pr5*, *Tnf*, *Ifng*, *Gzmb* and *Prf1* as an effector and effector memory score, both as described by Pauken et al. [Pauken et al.]. A cytotoxic score combined the known cytotoxic genes *Prf1*, *Ifng*, *Nkg7*, *Gzmb*, *Gzma*, *Gzmm*, *Klrk1*, *Klrb1a*, *Klrb1c*, *Klrd1*, *Ctsw* and *Cst7*. The migration score was taken from Hashimoto et al. and comprised *Cxcr3*, *S1pr1*, *Klf2*, *Itgb1*, *Cd44* and *Ly6c2* [Hashimoto et al.].

Scores were compared across metadata categories (matching category, tissue, treatment, recurrence) or combinations thereof using boxplots, and compared statistically using the Wilcoxon rank sum test, chosen because module scores are bounded and not normally distributed. In addition, the standard deviation of functional scores within a treatment was used as a measure of how treatment affects the functional diversity of tm T cells. To determine how tissue and treatment affect T cell subtypes, proportional Seurat cluster distributions were plotted with ggplot and compared against IgG treatment using a chi-squared test. Finally, to understand how relative function changes within a clone across tissues and in the presence or absence of antigen, functional scores were averaged per tissue per clone, each clonal average was plotted with geom_point, and matching clones were connected by a line and colored by tissue. To compare functional scores and gene changes across all mice, tissues and scores, each score was transformed to a Z-score with a mean of zero and a standard deviation of one. Differential expression was performed with FindMarkers or FindAllMarkers to determine changes between clusters, tissues or treatments, with min.pct and logfc.threshold set to 0.25, and visualized with EnhancedVolcano using a pCutOff of 0.05 and an FCcutoff of 0.5.

### Single-cell RNA pseudotime analysis

Monocle3 was used to perform a pseudotime analysis to study persistent clone-specific and treatment-specific T cell trajectories. T cells were first filtered to include only persisting clones. A new UMAP was created in Seurat to classify T cell subtypes using functional annotations, differential expression with FindAllMarkers, and functional scores with AddModuleScore. The Seurat object along with its metadata was then converted to a cell_data_set object, which was preprocessed, aligned by mouse ID, and passed to reduce_dimensions as recommended for Monocle3. Cells were then clustered with cluster_cells using the Louvain method, and the trajectory was determined with learn_graph. The earliest principal node was set in the tdLN, and its placement was verified against known markers of less differentiated states so that the trajectory root reflected biology rather than an arbitrary graph vertex. The resulting pseudotime trajectory was compared against the previously annotated T cell subtypes, tissue of origin and functional scores.

---

**Supplementary Figure 1.**
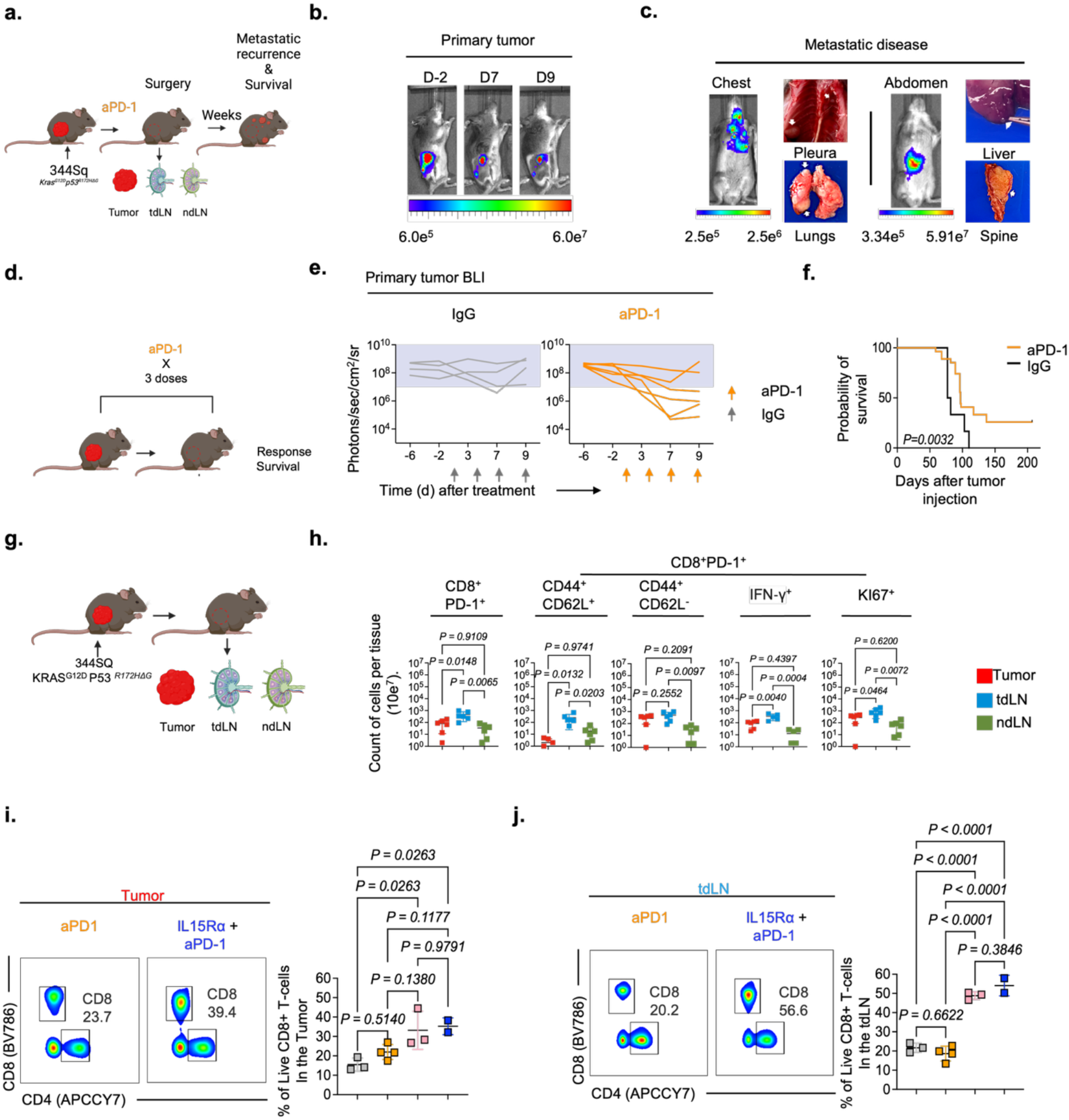
Surgical murine model displays natural tdLN CD8+ T cell responses when compared with the contralateral ntdLN. a. Schematic design. b. Representative bioluminescence (BLI) images of flank tumor burden of an aPD-1 responder mouse on days -2, 7 and 9 of treatment. c. Representative bioluminescence images and necropsy photographs of metastatic disease developing after primary tumor resection, showing common sites of metastasis. d. Experimental design using aPD-1. e. BLI plots of tumor growth kinetics during treatment with IgG control (gray) or aPD-1 (orange). f. Kaplan Meier survival plot of mice treated with aPD-1 or IgG. g. Experimental design for tumor, tumor-draining lymph node (tdLN) and non-tumor-draining lymph node (ntdLN) characterization. h. Flow cytometry quantification of CD8+ T cells in tumor, tdLN and ntdLN, showing counts of CD8+PD-1+ T cells and memory subpopulations determined by CD44 and CD62L expression; IFN-γ and Ki67 expression between tissues was also evaluated (6 biological replicates per tissue, n = 6). i, j. Representative flow cytometry plots and quantification of CD8+ T cells in the tumor and tumor-draining lymph node from mice treated with IgG (gray), aPD-1 (orange), IL-15 superagonist (pink) or aPD-1 + IL-15 superagonist (blue) after treatment. Statistics used were one-way analysis of variance (ANOVA) (h, i, j) and log-rank (Mantel Cox) test (f).

**Supplementary Figure 2.**
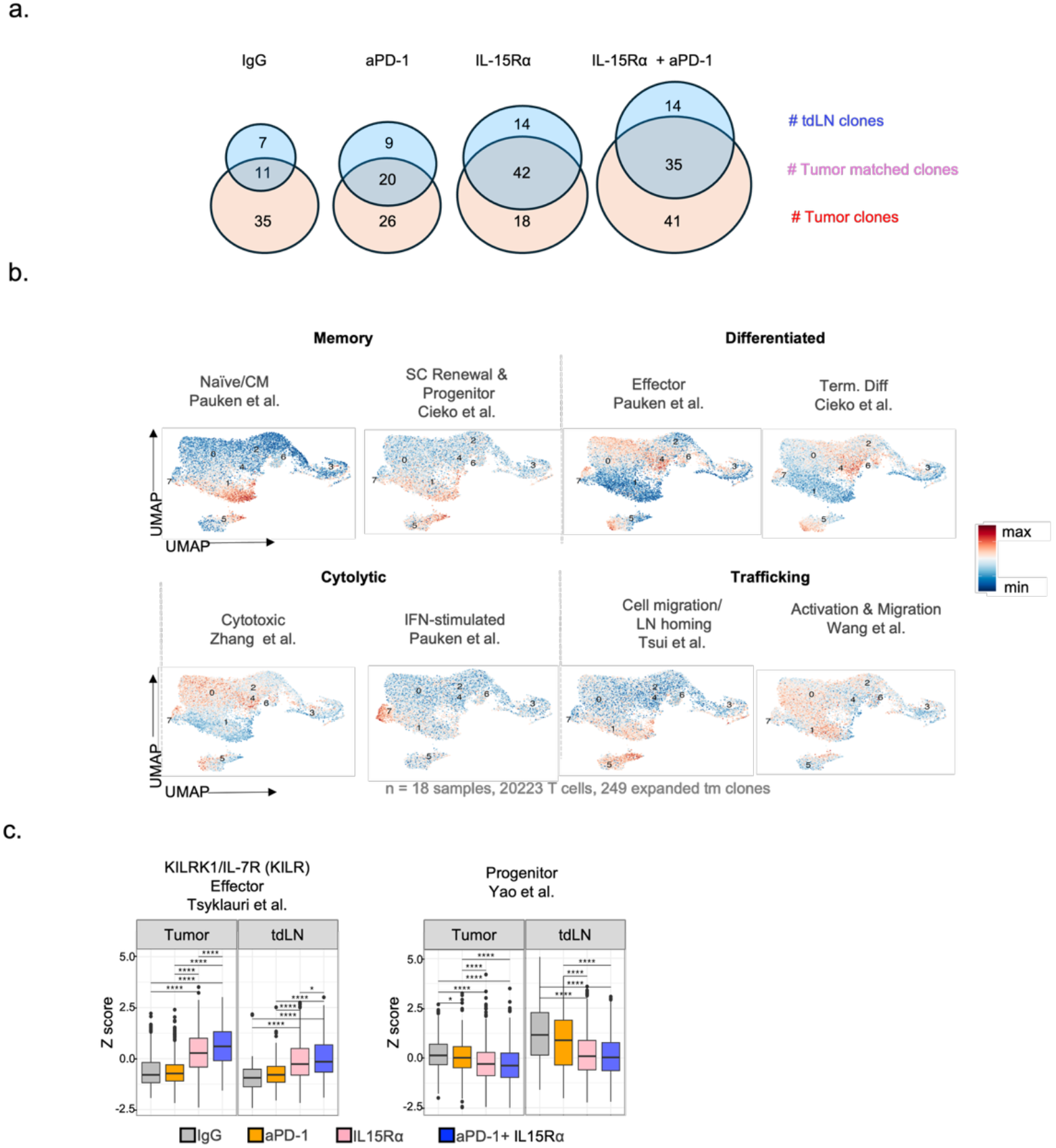
Single cell sequencing shows differences in CD8+ T cell repertoire dynamics induced by treatment. a. Venn diagram showing the number of clones from tumor (red), tdLN (blue) and overlapping clones between both tissues (tumor-matched clones) from the different treatments. b. UMAP showing relative expression of memory, differentiated, cytolytic and trafficking signatures. c. Z score for cytotoxic and KLRK/Il7r effector and progenitor signatures. Statistical analyses were t-test (c).

**Supplementary Figure 3.**
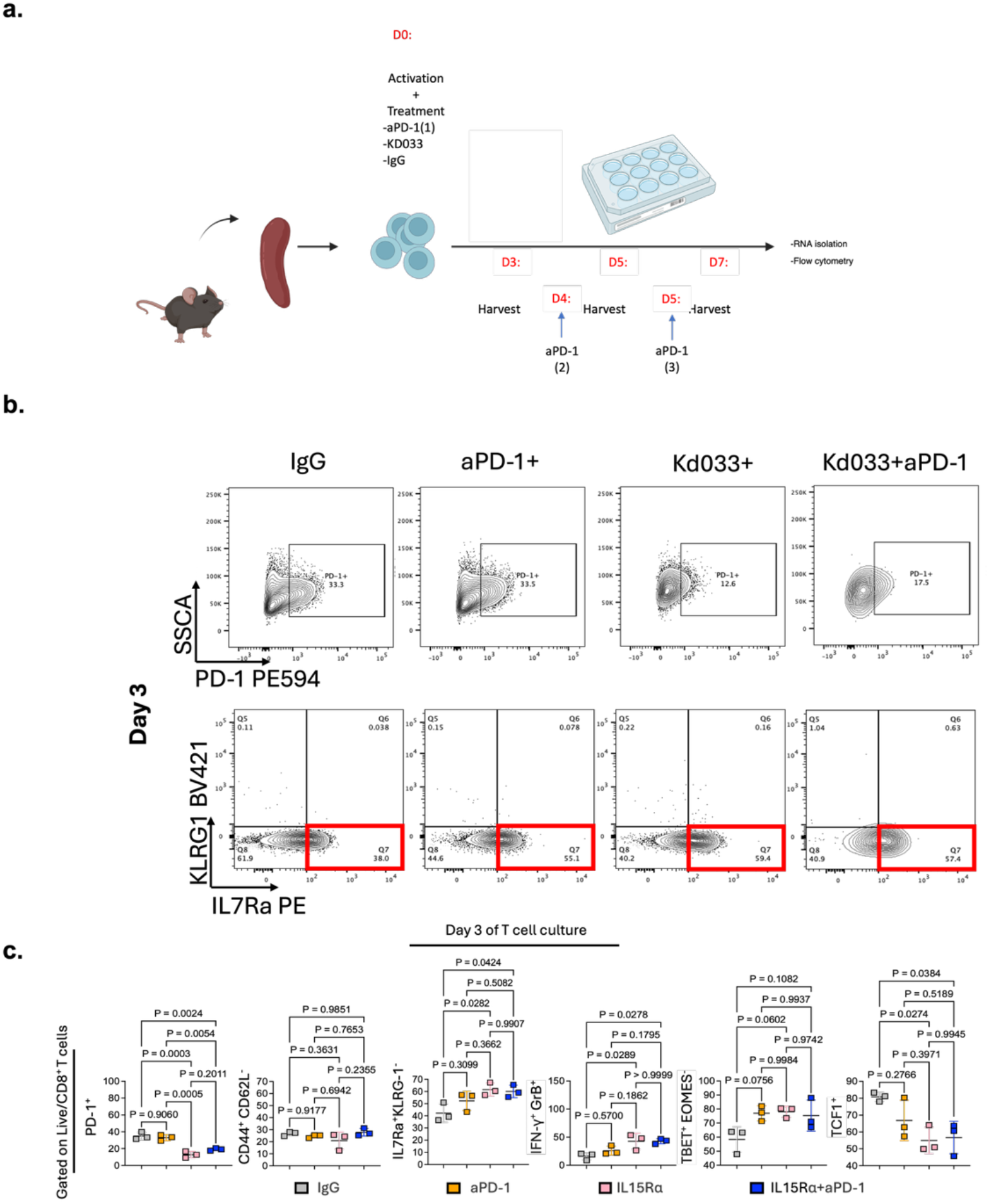
In vitro culture system for KD033 phenotypic characterization recapitulates the in vivo phenotype. a. Schematic design. b. Representative flow cytometry plots of CD8+ T cells at day 3 of in vitro culture. c. Flow cytometry quantification of CD8+ T cells treated with IgG (gray), aPD-1 (orange), IL-15 superagonist (pink) or aPD-1 + IL-15 superagonist (blue) at day 3 of culture. Statistics used were one-way analysis of variance (ANOVA) (c).

**Supplementary Figure 4.**
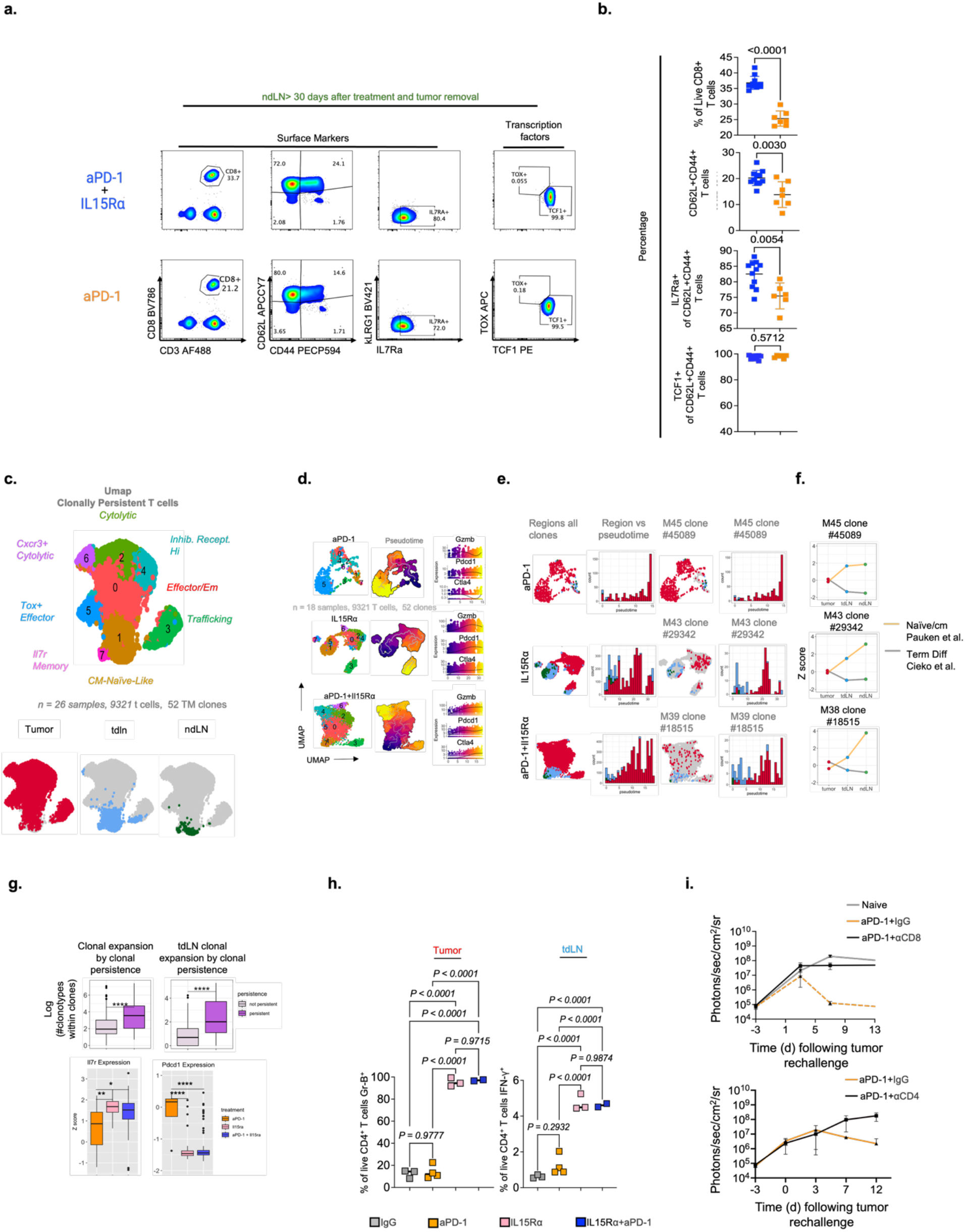
Systemic memory phenotype and pseudotime analysis show differences in CD8+ T cell fates. a. UMAP of clonally persistent CD8 T cells (n = 26 samples, 9,321 T cells, 52 tumor-matched clones), with transcriptional clusters labeled by functional annotation (CM/naïve-like, *Il7r* memory, trafficking, effector/effector memory, inhibitory receptor, cytolytic, Cxcr3+ cytolytic, Tox+ effector), and the same UMAP split by tissue of origin showing tumor (red), tdLN (blue) and ntdLN (green). b. UMAP localization, pseudotime, and *Gzmb*, *Pdcd1* and *Ctla4* expression of persistent clones for each treatment group: aPD-1, IL-15 superagonist, and aPD-1 + IL-15 superagonist (n = 18 samples, 9,321 T cells, 52 clones). c. Localization of all persistent clones by tissue of origin (left), distribution of tissue of origin across pseudotime (second), and UMAP localization with the corresponding pseudotime distribution for the largest clone in each treatment group: M45 clone #45089 (aPD-1), M43 clone #29342 (IL-15 superagonist) and M39 clone #18515 (aPD-1 + IL-15 superagonist). d. Average Z score for the naïve/central memory (Pauken et al.) and terminal differentiation (Ciecko et al.) signatures in the tumor, tdLN and ntdLN for the largest clone per treatment group. e. Clonal expansion, expressed as log clone size, compared between non-persistent and persistent clones across all tissues (left) and within the tdLN (right), and Z scores for *Il7r* and *Pdcd1* expression compared between aPD-1, IL-15 superagonist, and aPD-1 + IL-15 superagonist treated mice. f. KILR effector signature (Tsyklauri et al.) Z scores within the memory subclusters, compared between non-persistent and persistent clones for each treatment group (left), and receiver operating characteristic curves for discrimination of persistent clones by this signature in lymph node (AUC 0.766) and tumor (AUC 0.628) samples (right). g. Flow cytometry quantification of granzyme B expressing CD4+ T cells in the tumor and IFN-γ expressing CD4+ T cells in the tdLN after treatment with IgG, aPD-1, IL-15 superagonist, or aPD-1 + IL-15 superagonist. P values are shown on the plots. h. BLI plots of tumor burden after tail-vein rechallenge of aPD-1 treated, recurrence-free mice receiving IgG or CD8-depleting antibody, with naïve age-matched mice as controls (top), and IgG or CD4-depleting antibody (bottom). Box plots show median and interquartile range with whiskers to 1.5 times the interquartile range. *p < 0.05, **p < 0.01, ****p < 0.0001. Statistics used were Wilcoxon rank sum test for the two-group comparisons in e and f, Kruskal-Wallis with pairwise Wilcoxon testing for the three-group comparisons in e, and one-way analysis of variance (ANOVA) in g.

## References

1. Pu, X., et al., Immune Checkpoint Inhibitor Based Therapy as the First Line Treatment for Advanced Non Small Cell Lung Cancer: Efficacy, Challenges, and Future Perspectives. Thoracic Cancer. 16(12).

2. Ahmadzadeh, M., et al., Tumor antigen–specific CD8 T cells infiltrating the tumor express high levels of PD-1 and are functionally impaired. Blood, 2009. 114(8): p. 1537–1544.

3. Barber, D.L., et al., Restoring function in exhausted CD8 T cells during chronic viral infection. Nature, 2006. 439(7077): p. 682-687.

4. Zehn, D., et al., ‘Stem-like’ precursors are the fount to sustain persistent CD8+ T cell responses. Nature Immunology, 2022. 23(6): p. 836–847.

5. Im, S.J., et al., Defining CD8+ T cells that provide the proliferative burst after PD-1 therapy. Nature, 2016. 537(7620): p. 417-421.

6. Kallies, A., D. Zehn, and D.T. Utzschneider, Precursor exhausted T cells: key to successful immunotherapy? Nature Reviews Immunology, 2020. 20(2): p. 128–136.

7. Nose, Y., et al., PD-1 antibody-bound progenitor-exhausted CD8+ T cells in lymph nodes boost PD-1-blockade anti-tumor immunity in gastrointestinal cancer. Nature Communications, 2026. 17(1).

8. Hor, J.L., et al., Inhibitory PD-1 axis maintains high-avidity stem-like CD8+ T cells. Nature, 2026. 649(8095): p. 194-204.

9. Schenkel, J.M., et al., Conventional type I dendritic cells maintain a reservoir of proliferative tumor-antigen specific TCF-1+ CD8+ T cells in tumor-draining lymph nodes. Immunity, 2021. 54(10).

10. Li, Z., et al., In vivo labeling reveals continuous trafficking of TCF-1+ T cells between tumor and lymphoid tissue. Journal of Experimental Medicine, 2022. 219(6).

11. Rahim, M.K., et al., Dynamic CD8+ T cell responses to cancer immunotherapy in human regional lymph nodes are disrupted in metastatic lymph nodes. Cell, 2023. 186(6): p. 1127–1143.e18.

12. Utzschneider, D.T., et al., T Cell Factor 1-Expressing Memory-like CD8+ T Cells Sustain the Immune Response to Chronic Viral Infections. Immunity, 2016. 45(2): p. 415–427.

13. Miller, B.C., et al., Subsets of exhausted CD8+ T cells differentially mediate tumor control and respond to checkpoint blockade. Nature Immunology, 2019. 20(3): p. 326–336.

14. Forde, P.M., et al., Overall Survival with Neoadjuvant Nivolumab plus Chemotherapy in Lung Cancer. New England Journal of Medicine, 2025.

15. Rochman, Y., R. Spolski, and W.J. Leonard, New insights into the regulation of T cells by γc family cytokines. Nature Reviews Immunology, 2009. 9(7): p. 480–490.

16. Yang, Y. and A. Lundqvist, Immunomodulatory Effects of IL-2 and IL-15; Implications for Cancer Immunotherapy. Cancers, 2020. 12(12): p. 3586.

17. Atkins, M.B., et al., High-Dose Recombinant Interleukin 2 Therapy for Patients With Metastatic Melanoma: Analysis of 270 Patients Treated Between 1985 and 1993. J Clin Oncol 1999. 17.

18. Yang, J.C. and S.A. Rosenberg, An ongoing prospective randomized comparison of interleukin-2 regimens for the treatment of metastatic renal cell cancer. Cancer J Sci Am, 1997. 1:S79-84.

19. Hashimoto, M., et al., PD-1 combination therapy with IL-2 modifies CD8+ T cell exhaustion program. Nature, 2022. 610(7930): p. 173-181.

20. Hashimoto, M., S.S. Ramalingam, and R. Ahmed, Harnessing CD8 T cell responses using PD-1–IL-2 combination therapy. Trends in Cancer, 2024. 10(4): p. 332–346.

21. Waldmann, T.A., The Shared and Contrasting Roles of IL2 and IL15 in the Life and Death of Normal and Neoplastic Lymphocytes: Implications for Cancer Therapy. Cancer Immunology Research, 2015. 3(3): p. 219–227.

22. Melchionda, F., et al., Adjuvant IL-7 or IL-15 overcomes immunodominance and improves survival of the CD8+ memory cell pool. Journal of Clinical Investigation, 2005. 115(5): p. 1177–1187.

23. Lee, J., et al., IL-15 promotes self-renewal of progenitor exhausted CD8 T cells during persistent antigenic stimulation. Frontiers in Immunology, 2023. 14.

24. Zhang, X., et al., Potent and Selective Stimulation of Memory-Phenotype CD8+ T Cells In Vivo by IL-15. Immunity, 1998. 8(5): p. 591–599.

25. Schluns, K.S., K.D. Klonowski, and L. Lefranc Ois, Transregulation of memory CD8 T-cell proliferation by IL-15Rα+ bone marrow–derived cells. Blood, 2004. 103(3): p. 988–994.

26. Herndler-Brandstetter, D., et al., KLRG1+ Effector CD8+ T Cells Lose KLRG1, Differentiate into All Memory T Cell Lineages, and Convey Enhanced Protective Immunity. Immunity, 2018. 48(4): p. 716–729.e8.

27. McManus, D.T., et al., An early precursor CD8+ T cell that adapts to acute or chronic viral infection. Nature, 2025. 640(8059): p. 772-781.

28. Boersma, B., H. Poinot, and A. Pommier, Stimulating the Antitumor Immune Response Using Immunocytokines: A Preclinical and Clinical Overview. Pharmaceutics, 2024. 16(8): p. 974.

29. Lee, H., S.-H. Park, and E.-C. Shin, IL-15 in T-Cell Responses and Immunopathogenesis. Immune Network, 2024. 24(1).

30. Dammeijer, F., et al., The PD-1/PD-L1-Checkpoint Restrains T cell Immunity in Tumor-Draining Lymph Nodes. Cancer Cell, 2020. 38(5): p. 685–700.e8.

31. Fransen, M.F., et al., Tumor-draining lymph nodes are pivotal in PD-1/PD-L1 checkpoint therapy. JCI Insight, 2018. 3(23).

32. Ruggiu, M., et al., Anti-PD-1 therapy triggers Tfh cell–dependent IL-4 release to boost CD8 T cell responses in tumor-draining lymph nodes. Journal of Experimental Medicine, 2024. 221(4).

33. Ho, W.J., et al., Multipanel mass cytometry reveals anti–PD-1 therapy–mediated B and T cell compartment remodeling in tumor-draining lymph nodes. JCI Insight, 2020. 5(2).

34. Martomo, S.A., et al., Single-Dose Anti–PD-L1/IL-15 Fusion Protein KD033 Generates Synergistic Antitumor Immunity with Robust Tumor-Immune Gene Signatures and Memory Responses. Molecular Cancer Therapeutics, 2021. 20(2): p. 347–356.

35. Gibbons, D.L., et al., Contextual extracellular cues promote tumor cell EMT and metastasis by regulating miR-200 family expression. Genes & Development, 2009. 23(18): p. 2140–2151.

36. Harrell, M.I., B.M. Iritani, and A. Ruddell, Lymph Node Mapping in the Mouse. j Immunol Methods, 2008. 332(1-2).

37. Pauken, K.E., et al., Single-cell analyses identify circulating anti-tumor CD8 T cells and markers for their enrichment. Journal of Experimental Medicine, 2021. 218(4).

38. Andreatta, M., et al., Interpretation of T cell states from single-cell transcriptomics data using reference atlases. Nature Communications, 2021. 12(1).

39. Ayers, M., et al., IFN-γ–related mRNA profile predicts clinical response to PD-1 blockade. Journal of Clinical Investigation, 2017. 127(8): p. 2930–2940.

40. Kaech, S.M., et al., Molecular and Functional Profiling of Memory CD8 T Cell Differentiation. Cell, 2002. 111(6): p. 837–851.

41. Doering, T.A., et al., Network Analysis Reveals Centrally Connected Genes and Pathways Involved in CD8+ T Cell Exhaustion versus Memory. Immunity, 2012. 37(6).

42. Micevic, G., et al., IL-7R licenses a population of epigenetically poised memory CD8 T cells with superior antitumor efficacy that are critical for melanoma memory. Proceedings of the National Academy of Sciences, 2023. 120(30).

43. Chihara, N., et al., Induction and transcriptional regulation of the co-inhibitory gene module in T cells. Nature, 2018. 558.

44. Aoki, H., et al., Clonal Spreading of Tumor-Infiltrating T Cells Underlies the Robust Antitumor Immune Responses. Cancer Immunology Research, 2023. 11(6).

45. Aoki, H., et al., TCR Repertoire Analysis Reveals Mobilization of Novel CD8+ T Cell Clones Into the Cancer-Immunity Cycle Following Anti-CD4 Antibody Administration. Frontiers in Immunology, 2019. 9.

46. Gao, H., et al., Advances in IL–15–Based Cancer Immunotherapy and Divergent Immunological Effects of IL–2 and IL–15 Signaling via the Shared IL–2Rβγ Receptor. Frontiers in Immunology, 2026. 17.

47. Jiang, S., et al., A robust multi-scale clustering framework for single-cell RNA-seq data analysis. Scientific Reports, 2025. 15(1).

48. Song, W.-M., et al., Unsupervised multi-scale clustering of single-cell transcriptomes to identify hierarchical structures of cell subtypes. 2024, Springer Science and Business Media LLC.

49. Song, W.-M., et al., Unsupervised multiscale clustering of single-cell transcriptomes to identify hierarchical structures of cell subtypes. GigaScience, 2025. 14.

50. Yao, C., et al., Single-cell RNA-seq reveals TOX as a key regulator of CD8+ T cell persistence in chronic infection. Nature Immunology, 2019. 20(7): p. 890–901.

51. Pol, J.G., et al., Effects of interleukin-2 in immunostimulation and immunosuppression. Journal of Experimental Medicine, 2020. 217(1): p. jem.20191247.

52. Saxton, R.A., et al., Structure-based decoupling of the pro- and anti-inflammatory functions of interleukin-10. Science, 2021. 371(6535): p. eabc8433.

53. Glassman, C.R., et al., Structural basis for IL-12 and IL-23 receptor sharing reveals a gateway for shaping actions on T versus NK cells. Cell, 2021. 184(4): p. 983–999.e24.

54. Tripathi, P., et al., STAT5 Is Critical To Maintain Effector CD8+ T Cell Responses. The Journal of Immunology, 2010. 185(4).

55. Hand, T.W., et al., Differential effects of STAT5 and PI3K/AKT signaling on effector and memory CD8 T-cell survival. Proceedings of the National Academy of Sciences, 2010. 107(38): p. 16601–16606.

56. Kemp, R.A., et al., Evidence of STAT5 dependent and independent routes to CD8 memory formation and a preferential role for IL-7 over IL-15 in STAT5 activation. Immunol Cell Biol..

57. Wang, G., et al., STAT5 interferes with PD-1 transcriptional activation and affects CD8+ T-cell sensitivity to PD-1-dependent immunoregulation. International Immunology, 2021. 33(11).

58. Ristin, S., et al., Asymmetry and redundancy of STAT5 paralogs across CD8+ T cell differentiation states. Communications Biology, 2026. 9(1).

59. Enamorado, M., et al., Enhanced anti-tumour immunity requires the interplay between resident and circulating memory CD8+ T cells. Nature Communications, 2017. 8(1).

60. Amaria, R.N., et al., Regulatable membrane-bound IL-15 engineered TIL cell therapy for immune checkpoint inhibitor-resistant advanced melanoma: pooled analysis of phase 1 and 1/2 trials. Nature Medicine, 2026.

61. A., R., et al., IL-2-independent expansion, persistence, and antitumor activity in TIL expressing regulatable membrane-bound IL-15. Molecular Therapy, 2025. 33(8).

62. Rodriguez, G.E., et al., Rewiring STAT signaling from the cell surface with Trikine immunotherapeutics. Science, 2026. 392(6799).

63. Luke, J.J., et al., Phase I dose escalation of KD033, a PDL1-IL15 bispecific molecule, in advanced solid tumors. JOURNAL OF CLINICAL ONCOLOGY, 2021. 39.

64. Chamie, K., et al., Final clinical results of pivotal trial of IL-15RαFc superagonist N-803 with BCG in BCG-unresponsive CIS and papillary nonmuscle-invasive bladder cancer (NMIBC). Journal of Clinical Oncology 2022. 40.

65. Markowitz, G.J., et al., Immune reprogramming via PD-1 inhibition enhances early-stage lung cancer survival. JCI Insight, 2018. 3(13).

